# A conserved SUMO-Ubiquitin pathway directed by RNF4/SLX5-SLX8 and PIAS4/SIZ1 drives proteasomal degradation of topoisomerase DNA-protein crosslinks

**DOI:** 10.1101/707661

**Authors:** Yilun Sun, Lisa M. Miller Jenkins, Yijun P. Su, Karin C. Nitiss, John L. Nitiss, Yves Pommier

## Abstract

Topoisomerase cleavage complexes (TOPccs) can be stalled physiologically and by the anticancer drugs camptothecins (TOP1 inhibitors) and etoposide (TOP2 inhibitor), yielding irreversible TOP DNA-protein crosslinks (TOP-DPCs). Here we elucidate how TOP-DPCs are degraded via the SUMO-ubiquitin (Ub) pathway. We show that in human cells, TOP-DPCs are promptly and sequentially conjugated by SUMO-2/3, SUMO-1 and Ub. SUMOylation is catalyzed by the SUMO ligase PIAS4, which forms a complex with both TOP1 and TOP2α and β. RNF4 acts as the SUMO-targeted ubiquitin ligase (STUbL) for both TOP1- and TOP2-DPCs in a SUMO-dependent but replication/transcription-independent manner. This SUMO-Ub pathway is conserved in yeast with Siz1 the ortholog of PIAS4 and Slx5-Slx8 the ortholog of RNF4. Our study reveals a conserved SUMO-dependent ubiquitylation pathway for proteasomal degradation of both TOP1- and TOP2-DPCs and potentially for other DPCs.

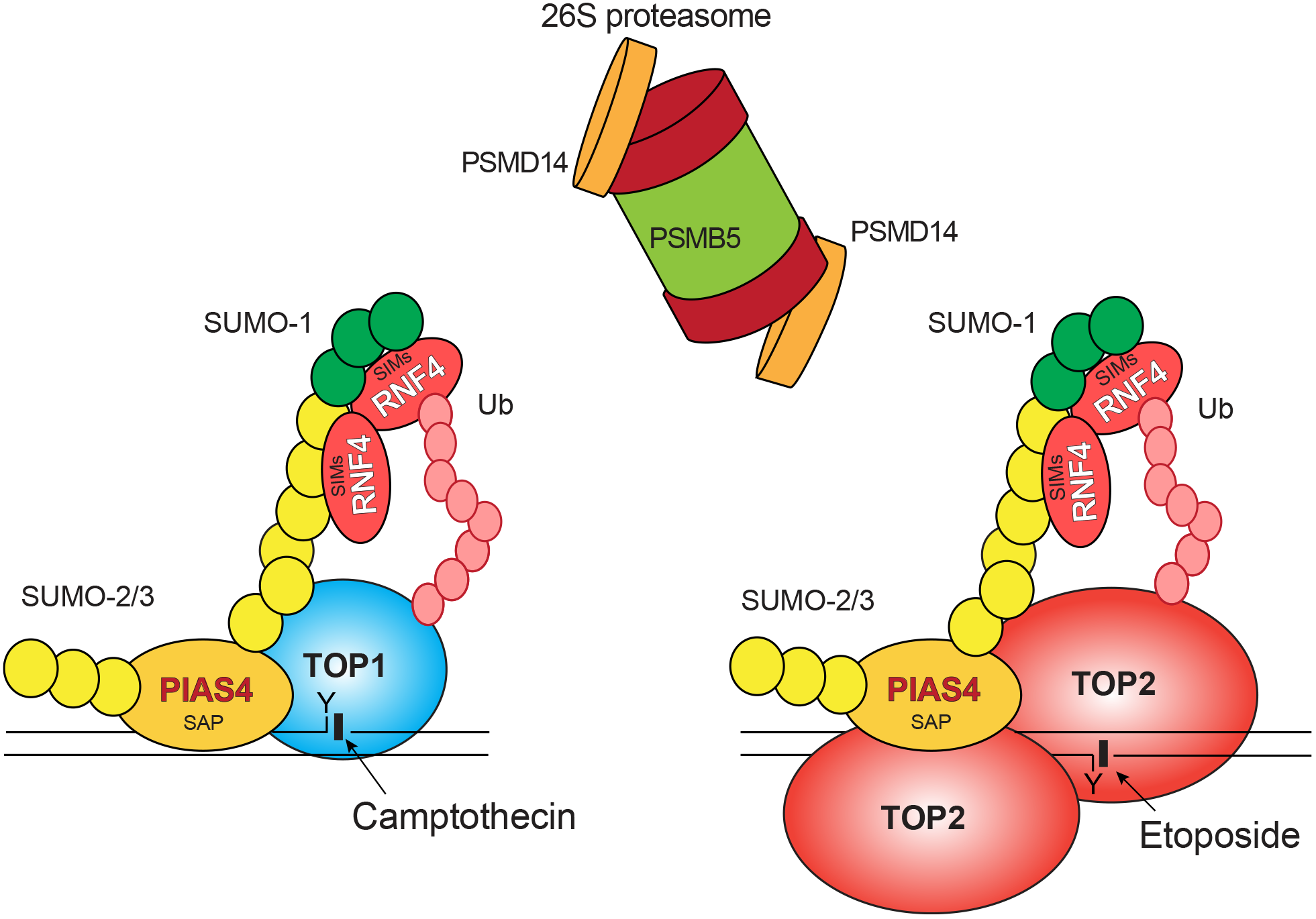

**In Brief:** Topoisomerase DNA-protein crosslinks (TOP-DPCs) are the therapeutic mechanism of clinical TOP inhibitors (camptothecin and etoposide). TOP-DPCs induce rapid and sequential conjugation of SUMO-2/3- SUMO-1 and ubiquitin catalyzed by activation of PIAS4 through its DNA-binding SAP domain and RNF4 through its SIM domains. This SUMO-ubiquitin cascade triggers proteasomal degradation of TOP-DPCs.

**HIGHLIGHTS:**

- Abortive topoisomerase I (TOP1) and II (TOP2) cleavage complexes resulting in DNA-protein crosslinks (TOP-DPCs) are rapidly and sequentially modified by SUMO-2/3, SUMO-1 and ubiquitin before their proteasomal degradation.
- PIAS4 SUMOylates TOP-DPCs via its DNA-binding SAP domain independently of DNA transactions and DNA damage responses.
- RNF4 ubiquitylates SUMOylated TOP-DPCs and drives their proteasomal degradation.
- TOP-DPC processing by the SUMO-Ub pathways is conserved in yeast and human cells.

## INTRODUCTION

DNA is constantly undergoing damage from endogenous and exogenous sources (Lindahl, 1993), which compromises genome integrity (Hoeijmakers, 2009). Endogenous DNA lesions are highly diverse, ranging from abasic sites, chemical adducts, single- and double-strand breaks (SSBs and DSBs) and DNA-protein crosslinks (DPCs) (Friedberg et al., 2014). Intrinsic defects in DNA repair systems leads to carcinogenesis, neurodegeneration, immunodeficiency and other deleterious consequences (Jackson and Bartek, 2009). Repair mechanisms specialized for DPCs are a current focus of investigation (Larsen et al., 2019; Stingele et al., 2017).

A ubiquitous mechanism for DPC formation is through the action of topoisomerases (TOPs). TOP1 and TOP2 cleave one or both strands of the DNA, respectively, through transient covalent linkage between their catalytic tyrosine residue and the phosphate group of the DNA backbone (3’-end for TOP1 and 5’-end for TOP2), leading to formation of TOP-DNA cleavage complexes (TOPccs). Upon re-ligation of the DNA, TOPs are released from the DNA (Pommier et al., 2016; Vos et al., 2011). Failure in the re-ligation step results in persistent TOPccs (TOP-DPCs) that interfere with DNA metabolisms upon collisions with DNA and RNA polymerases. Although TOP1 and TOP2 are distinct proteins in terms of their sizes, structures and catalytic mechanisms, TOP-DPCs can stem from topoisomerase malfunctions (Fertala et al., 2000; Pouliot et al., 1999), as well as preexisting DNA lesions such as abasic sites, oxidative damage, mismatches, ribonucleotide misincorporation and strand breaks (Pommier et al., 2016). TOP-DPCs are also induced therapeutically by the anticancer agents camptothecin (CPT) derivatives (TOP1 poisons) and etoposide (ETP), doxorubicin or amsacrine (TOP2 poisons).

TOP-DPC-specific repair mechanisms are generally categorized into two classes: nucleolytic cleavage and proteolytic degradation. The first class is exemplified by tyrosine-DNA phosphodiesterases (TDPs) that hydrolyze the covalent tyrosine-DNA bond, with TDP1 excising TOP1-DPCs (Pouliot et al., 1999) and TDP2 excising TOP2-DPCs in higher eukaryotes (Cortes Ledesma et al., 2009). In additions to TDPs, endonucleases such as Mre11 and XPF have been reported to play a key role in removing both TOP-DPCs presumably by incising the DNA adjacent to the TOP-DPC (Aparicio et al., 2016; Pommier et al., 2016; Sasanuma et al., 2018). Proteolytic degradation is required for TDP activities and earlier work showed that the ubiquitin (Ub)-proteasome pathway (UPP) plays an important role in proteolyzing TOP-DPCs (Desai et al., 1997; Mao et al., 2001). Yet, the detailed molecular mechanisms by which the UPP targets TOP-DPCs are still poorly understood. In addition, small ubiquitin-like modifiers (SUMO) 1 and 2/3 have been implicated in TOP-DPC repair (Agostinho et al., 2008; Kanagasabai et al., 2009; Mao et al., 2000a; Mao et al., 2000b), but the functions of these modifications still remain elusive.

SUMOylation has been linked to ubiquitylation, as polymeric SUMO chains can serve as mark for recognition by SUMO-targeted ubiquitin ligases (STUbLs) that attach Ub moieties on poly-SUMOylated substrates to promote their proteasomal degradation (Poulsen et al., 2013; Tatham et al., 2008; van Hagen et al., 2010). SUMO-Ub pathways orchestrate the proteolytic turnover of DNA damage response (DDR) and repair enzymes to facilitate DNA repair in mammalian cells (Jackson and Durocher, 2013; Ulrich, 2012). The Siz/PIAS RING family SUMO ligases PIAS1 and PIAS4 mediate SUMOylation as an early response to DNA damage by modifying DDR proteins including MDC1, RPA and 53BP1 to enable their recruitment to DNA damage sites (Galanty et al., 2009). Among STUbLs, RNF4 is implicated in the ubiquitylation and proteolysis of SUMOylated substrates to ensure DSB repair (Galanty et al., 2012; Vyas et al., 2013; Yin et al., 2012). In *Saccharomyces cerevisiae*, the PIAS family orthologs Siz1 and Siz2 as well as the RNF4 ortholog Slx5-Slx8 heterocomplex engage in DNA repair to ensure genome stability (Heideker et al., 2011; Jalal et al., 2017; Zhang et al., 2006b).

Here we dissect the involvement of SUMO, Ub and the proteasome in the repair of TOP-DPCs. We elucidate the SUMO-Ub axis in yeast with Siz1 responsible for TOP1- and TOP2-DPC SUMOylation and ubiquitylation by Slx5-Slx8. Using a novel assay for detection of ubiquitylated and SUMOylated TOP-DPCs (DUST), we show for the first time that TOP1- and TOP2-DPCs are promptly modified by SUMO-2/3, SUMO-1 and Ub in a sequential order in human cells. We identify human PIAS4 as the major SUMO ligase for both TOP1- and TOP2-DPCs. We also show that RNF4 is the STUbL for TOP-DPCs in human cells, which drives their proteasomal degradation.

## RESULTS

### SUMO-Ub Axis and Proteasome Repair of TOP-DPC in Yeast

The UPP pathway is conserved in eukaryotes and ubiquitin ligases that specifically recognize SUMOylated proteins have been discovered in species ranging from yeasts to humans (Sriramachandran and Dohmen, 2014). To investigate whether UPP is involved in the processing of TOP-DPCs, Western blotting (WB) was carried out in the yeast *S. cerevisiae* strain YMM10 overexpressing HA-tagged yeast TOP1 or TOP2. The results revealed that CPT and ETP induce partial loss of cellular TOP1 and TOP2 (Figure S1A, B), which was blocked by the proteasome inhibitor MG132, indicating TOP proteasomal degradation upon TOP-DPC formation. Clonogenic assays showed that MG132 rendered TOP1- and TOP2-overexpressing cells hypersensitive to CPT and ETP, respectively (Figure S1C, D). These results demonstrate the proteasome as a conserved pathway for the repair of TOP-induced DNA damage.

We next examined whether yeast TOPs are modified by Ub and SUMO upon DPC formation. HA-antibody immunoprecipitation (IP) showed that both TOP1 and TOP2 are conjugated with polymeric Ub and smt3 (yeast SUMO-1) at steady-state, and that addition of CPT or ETP enhanced formation of smt3- and Ub-TOP conjugates (Figure 1A, B). As IP of cellular lysates cannot distinguish between free TOPs and TOP-DPCs, we developed an approach to detect DPC post translational modifications (PTMs). We modified and adapted the classical *in vivo* complex of enzyme (ICE) assay (Anand et al., 2018; Subramanian et al., 1995). In brief, we digested the isolated TOP-DPCs with micrococcal nuclease and resolved the DPCs by SDS-PAGE gel electrophoresis, followed by immunoblotting (IB) to assess TOP-DPC and their PTMs using specific antibodies. Both TOP1- and TOP2-DPCs as well as their ubiquitylation and SUMOylation were detected in the absence of TOP inhibitors, presumably as a result of overexpression of TOPs (Figure 1C, D). Treatment with CPT and ETP further induced TOP-DPCs and MG132 stimulated TOP-DPC accumulation, indicating a role of the proteasome in eliminating TOP-DPCs. Concurrently, TOP inhibitor treatments increased the levels of smt3- and Ub-TOP-DPCs, which were further increased by proteasome inhibition.

**Figure 1.**
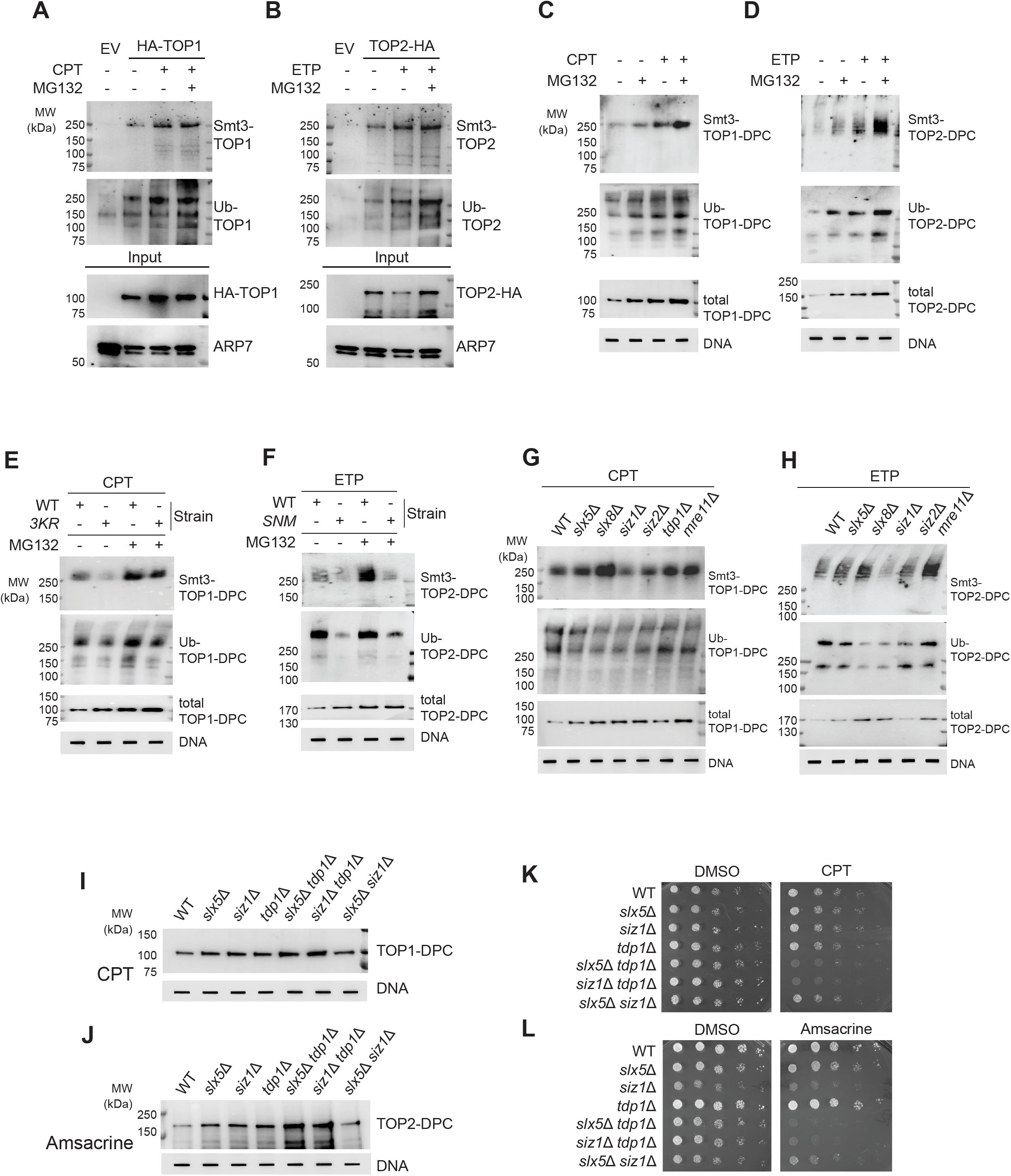
SUMO-Ubiquitin Pathway and Proteasome Repair of TOP-DPCs in Yeast. (A) YMM10 cells carrying empty vector (EV) or pYX112 HA-TOP1 plasmid were treated with DMSO or CPT (20 μg/ml) ± MG132 (pre-treatments at 10 μM for 1 h) for 30 min, followed by HA-IP and immunoblotting (IB) with indicated antibodies. (B) YMM10 cells carrying the pDED1 TOP2-3×HA plasmid were treated with DMSO, ETP (200 μg/ml) ± MG132 for 30 min, followed by HA-IP and IB. (C) ICE assays in YMM10 cells overexpressing HA-TOP1 after treatment with DMSO, MG132, CPT (20 μg/ml) or CPT + MG132 for 1 h. smt3-, Ub and total TOP1-DPCs were detected with indicated antibodies. Undigested ICE samples were probed with anti-dsDNA antibody as loading control. (D) ICE assay in YMM10 cells overexpressing TOP2-HA after treatment with DMSO, MG132, ETP (200 μg/ml) or ETP + MG132 for 1 h. (E) YMM10 cells overexpressing HA-TOP1 WT or HA-*top1 K65,91,92R* were exposed to 20 μg/ml CPT ± MG132 for 1 h, followed by ICE assay with indicated antibodies. YMM10 cells overexpressing TOP2-HA WT or *top2 SNM*-HA were exposed to 200 μg/ml ETP ± MG132 for 1 h, followed by ICE assay. (G) After exposure to 20 μg/ml CPT for 1 h, levels of smt3-, Ub- and total TOP1-DPCs in the indicated BY4741 strains overexpressing HA-TOP1 were determined by ICE assay. (H) After exposure to 200 μg/ml ETP for 1 h, levels of smt3-, Ub- and total TOP2-DPCs in the indicated BY4741 strains overexpressing TOP2-HA were determined by ICE assay. (I) After exposure to 20 μg/ml CPT for 1 h, total TOP1-DPCs in the indicated BY4741 strains overexpressing HA-TOP1 were detected by ICE assay. (J) After exposure to 200 μg/ml amsacrine for 1 h, total TOP2-DPCs in the indicated strains overexpressing TOP2-HA were detected by ICE assay. (K) Liquid culture of the indicated BY4741 overexpressing HA-TOP1 were plated on DMSO or CPT (0.5 μg/ml) containing SC-URA plates for spot test. (L) Liquid culture of the indicated BY4741 strains overexpressing TOP2-HA were plated on DMSO or amsacrine (20 μg/ml) as in panel (K).

To investigate whether TOP-DPC SUMOylation plays a role in TOP-DPC ubiquitylation, we transformed YMM10 strain with the *top1 K65, 91, 92R* (*top1-3KR*) (Chen et al., 2007) or *top2 Sumo No More* (*top2-SNM*) (Bachant et al., 2002) constructs, both of which comprise mutations disrupting their respective SUMO consensus sites. Both the *top1K65,91, 92R* allele and *top2 SNM* allele showed a substantial decrease of SUMO-TOP-DPCs compared with the strains harboring the wild-type (WT) plasmids (Figure 1E, F, Figure S1E, F). Notably, deficiency in SUMOylation brought about marked reduction in both TOP1- and 2-DPC ubiquitylation, suggesting that SUMOylation licenses TOP-DPC for ubiquitylation in yeast.

### Siz1 and Slx5/Slx8 are SUMO ligase and STUbL for TOP-DPCs in Yeast

To identify the SUMO ligase(s) and STUbL(s) for TOP-DPC, we performed ICE assays in yeast strains lacking SUMO ligases and STUbLs involved in DNA damage repair in yeast. Deletion of the genes encoding the STUbL Slx5-Slx8 complex decreased TOP-DPC ubiquitylation (Figure 1g, h). Comparing *slx5* null and *slx8* null strains, *slx8*Δ displayed lower Ub-TOP-DPC levels than the slx5Δ strains (Figure 1G, H), consistent with a study showing that Slx8 rather than Slx5 mediates dsDNA binding activity of the complex (Yang et al., 2006a). Also, the *slx5*Δ and *slx8*Δ strains both showed elevated TOP-DPC SUMOylation, suggesting a role of Slx5-Slx8 in processing SUMOylated TOP-DPCs. Deletion of gene encoding SUMO ligase SIZ1 but not SIZ2 substantially reduced not only SUMOylation of TOP-DPCs but also their ubiquitylation (Figure 1G, H, Figure S1G, H), supporting the conclusion that SUMOylation prompts TOP-DPC ubiquitylation.

Because Tdp1 is a critical enzyme for excising TOP-DPCs, we assessed its relationship with Siz1, Slx5 and Tdp1. *slx5*Δ *tdp1*Δ and *siz1*Δ t*dp1*Δ double-mutants exhibited higher TOP-DPC levels than those in the respective single mutants (Figure 1I, J, Figure S1I, J). TOP-DPC levels were comparable in *slx5*Δ *siz1*Δ double-mutant and in *slx5*Δ and *siz1*Δ single-mutants, suggesting the epistasis of Siz1 and Slx5 in TOP-DPC repair. Next, we compared the sensitivity of the mutants to TOP inhibitors by spotting cultures onto plates containing CPT or TOP2 inhibitor amsacrine (Figure 1k, l). Substantially reduced growth of *slx5*Δ *tdp1*Δ and *siz1*Δ t*dp1*Δ double-mutant strains can be seen on the drug plates (Figure 1K, L). Taken together, these observations indicate redundant roles of Siz1-Slx5 and Tdp1 in removing TOP-DPCs in yeast.

### SUMO-, Ub- and Proteasome-dependent processing of TOP-DPCs in human cells

We next sought to examine the UPP-dependent repair of TOP-DPCs in human cells. First, we assessed the proteasomal degradation of TOP1, TOP2α and TOP2β in CPT- or ETP-treated human embryonic kidney HEK293 cells in the presence of MG132 and the Ub-activating enzyme (UAE) inhibitor TAK-243 (Hyer et al., 2018). CPT or ETP treatments for 4 h led to a marked reduction of TOP1, TOP2α and TOP2β levels (Figure 2A, B, lanes 1-4), and both MG-132 and TAK-243 blocked CPT-induced TOP1 degradation and ETP-induced TOP2α and β degradation (Figure 2A, B, lanes 5, 6), demonstrating UPP-dependent destruction of cellular TOPs in response to TOP poisoning. ICE assays showed that MG132 or TAK-243 increased CPT-induced TOP1-DPC and ETP-induced TOP2-DPC levels (Figure 2C-H), suggesting that UPP targets TOP-DPCs for the destruction.

**Figure 2.**
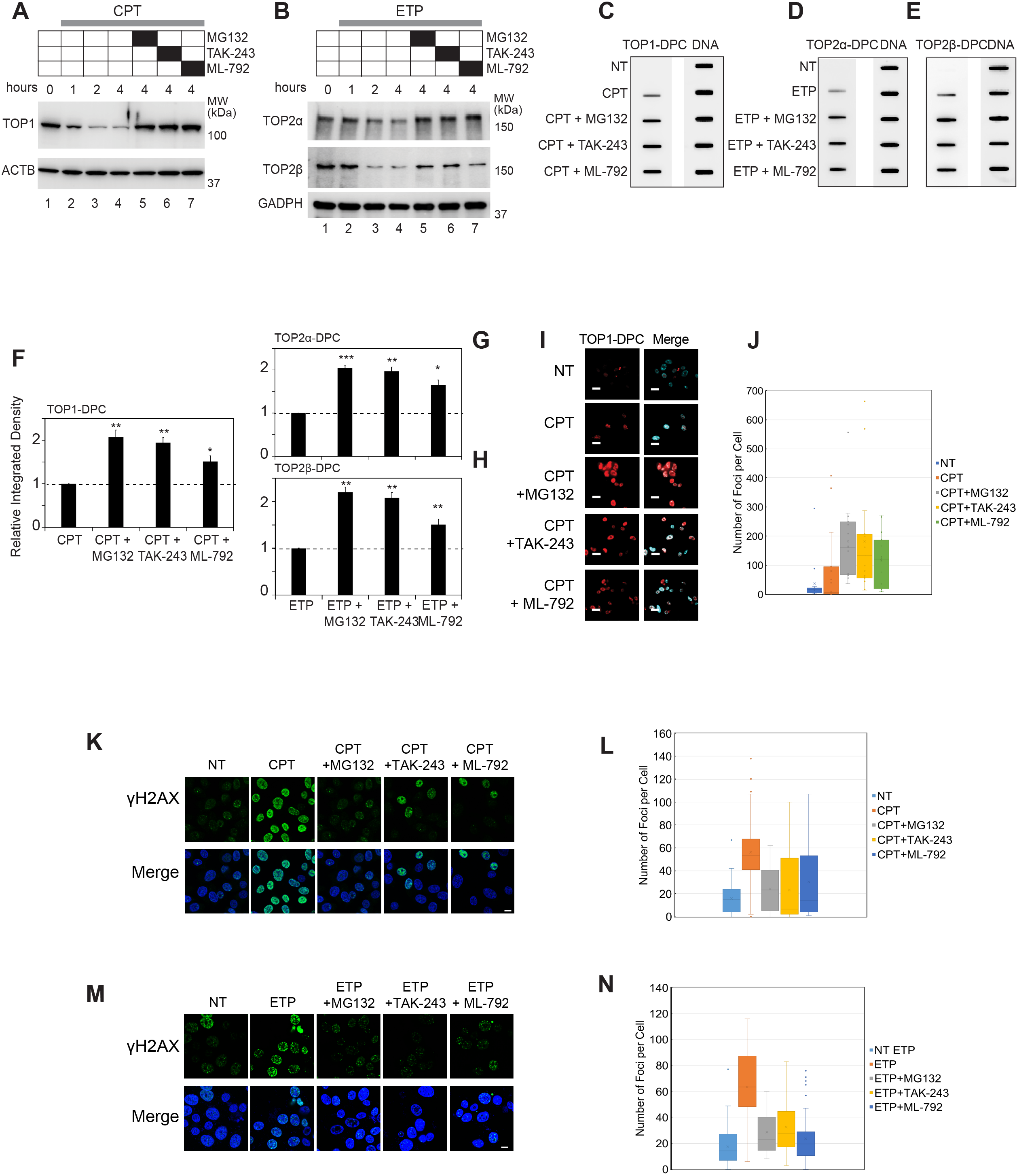
Proteasomal Degradation, Ubiquitylation and SUMOylation are Involved in the Resolution of TOP-DPCs in Human Cells. (A)&(B) HEK293 cells were treated with CPT (20 μM) or ETP (200 μM) as indicated. MG132 (10 μM) was added 1 h prior to CPT or ETP addition. The UAE inhibitor TAK-243 (10 μM) was added 2 h prior to CPT or ETP. The SAE inhibitor ML-792 (10 μM) was added 4 h prior to CPT or ETP. Cells were then lysed with the alkaline lysis procedure and immunoblotted as indicated. (C)-(E) HEK293 cells were treated with CPT (20 μM) or ETP (200 μM) as indicated. MG132 (10 μM) was added 1 h prior to CPT or ETP. The UAE inhibitor TAK-243 (10 μM) was added 2 h prior to CPT or ETP. The SAE inhibitor ML-792 (10 μM) was added 4 h prior to CPT or ETP. Cells were then subjected to ICE assay for immunodetection TOP-DPC using anti-TOP1, anti-TOP2α or anti-TOP2β antibodies. (F)-(H) Densitometric analyses comparing relative integrated densities of TOP1-DPC, TOP2α-DPC or TOP2β-DPC signals in independent experiments shown in panels (C)-(E). Integrated density of TOP-DPC signals of each group was normalized to that of cells treated with CPT or ETP alone. * p-values < 0.05; ** p-values < 0.01. (I) U2OS cells were treated with CPT (10 μM) for 1 h in the presence and absence of the indicated inhibitors. MG132 (10 μM) was added 1 h prior to CPT. TAK-243 (10 μM) was added 2 h prior to CPT. ML-792 (10 μM) was added 4 h prior to CPT. Cells were then subjected to immunofluorescence (IF) microscopy for detection of TOP1-DPC using anti-TOP1-DPC antibody as described under Materials and Methods. Scale bar represents 20 microns. (J) Quantitation of TOP1-DPC foci per cell of each treatment group from panel (I). (K) U2OS cells were treated with CPT (1 μM) for 1 h as indicated. MG132 (10 μM) was added 1 h prior to CPT, TAK-243 (10 μM) 2 h prior to CPT, and ML-792 (10 μM) 4 h prior to CPT. Cells were then subjected to IF for detection of γH2AX. Scale bar represents 20 microns. (L) Quantitation of γH2AX foci per cell of each treatment group from panel (K). (M) U2OS cells were treated with ETP (5 μM) for 1 h in the presence and absence of various inhibitors. MG132 (10 μM) was added 1 h prior to ETP addition. TAK-243 (10 μM) was added 2 h prior to ETP addition. ML-792 (10 μM) was added 4 h prior to ETP addition. Cells were then subjected to IF assay for detection of γH2AX. (N) Quantitation of γH2AX foci per cell of each treatment group from panel (M).

We then intended to establish a role of SUMOylation in TOP-DPC processing. Utilizing ML-792, an inhibitor of the SUMO-activating enzyme SAE (He et al., 2017), we found that inhibiting SUMOylation blocked drug-induced loss of TOPs (Figure 2A, B, lanes 7) while increasing TOP-DPCs (Figure S2C-H). Using an antibody that selectively recognizes TOP1-DPCs (Patel et al., 2016), immunofluorescence (IF) microscopy showed that inhibition of the proteasome, ubiquitylation or SUMOylation increased TOP1-DPCs (Figure 2I, J), corroborating the involvement of SUMOylation, ubiquitylation and proteasome in TOP1-DPC resolution.

Upon exposure to TOP inhibitors, induction of γH2AX, a marker for DSBs (Bonner et al., 2008) requires proteolysis, as evidenced by the observation that the MG132 blocks TOP inhibitor-induced γH2AX foci formation (Lin et al., 2009; Schellenberg et al., 2017; Zhang et al., 2006a). Pre-treatment with TAK-243 or ML-792 also reduced CPT- and ETP-induced γH2AX foci (Figure S2K-N), implying a role of SUMOylation and ubiquitylation in signaling downstream from TOP-DPCs.

### Rapid and Sequential SUMOylation and Ubiquitylation of TOP-DPCs in Human Cells

To investigate whether Ub and SUMO modify TOP-DPCs for repair in human cells, we performed pull-down experiments by transiently transfecting TOPs in HEK293 cells. His-tagged TOP1 was found conjugated with SUMO-2/3 and Ub at baseline, and CPT enhanced its SUMO-1, SUMO-2/3, and Ub modifications (Figure 3A). Proteasome inhibition by MG132 further stimulated TOP1 SUMOylation and ubiquitylation. Similarly, IP of FLAG-tagged TOP2 and TOP2 showed that TOP2 isozymes are SUMO-2/3-, SUMO-1- and Ub-conjugated at baseline, and that treatment with ETP enhanced their SUMOylation and ubiquitylation, which were increased upon co-treatment with MG132 (Figure 3B).

**Figure 3.**
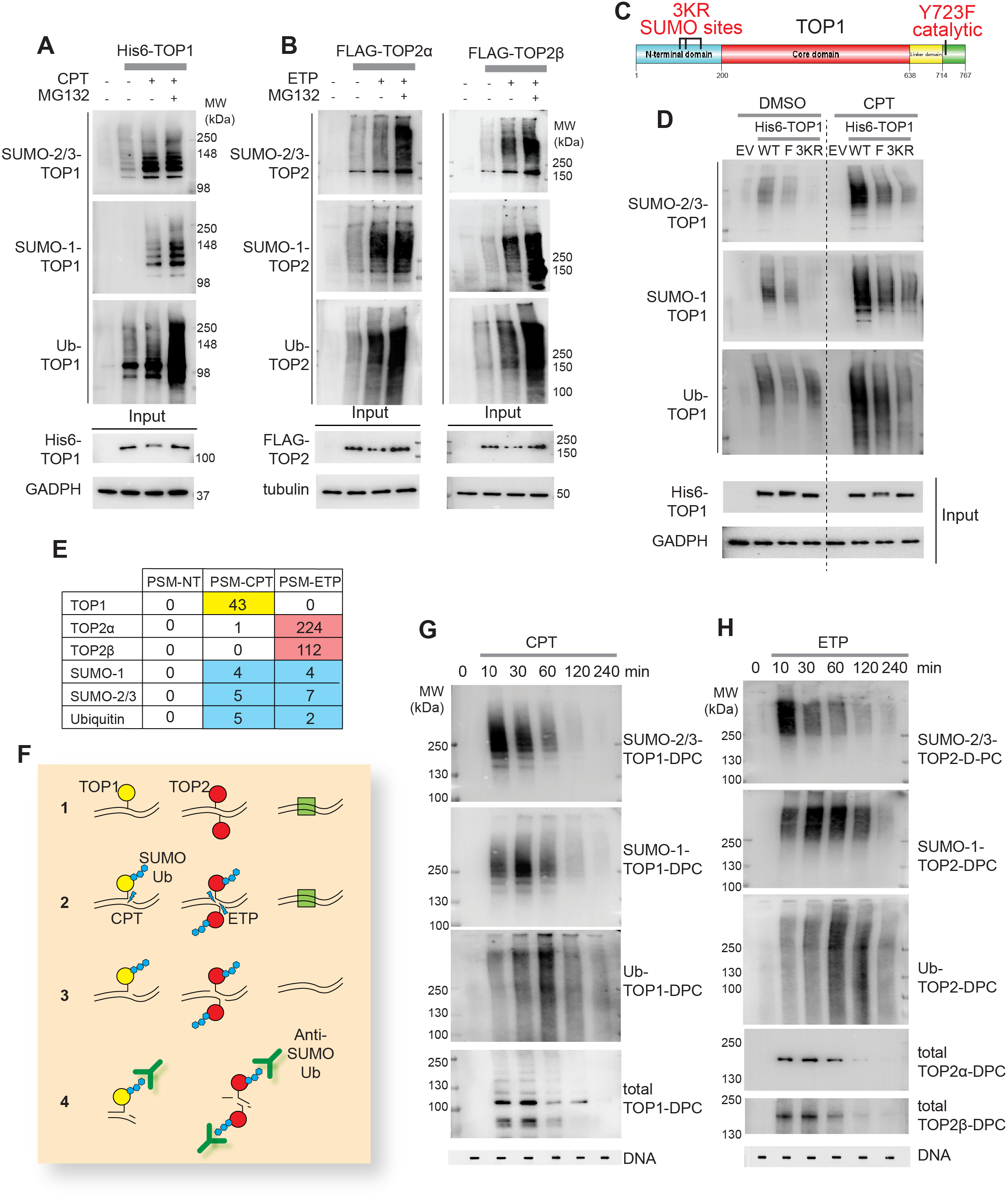
Rapid and Sequential SUMOylation and Ubiquitylation of TOP-DPCs in Human Cells. (A) HEK293 cells were transfected with the 6×His tagged TOP1 expression plasmid, followed by treatments with DMSO or CPT ± MG132 for 30 min. His-pull down samples and cell lysates (input) were subjected to IB with indicated antibodies. (B) HEK293 cells were transfected with the FLAG tagged TOP2α or TOP2β expression plasmids, followed by treatments with DMSO or ETP ± MG132 for 30 min. FLAG-IP samples and cell lysates were subjected to IB with indicated antibodies (C) Scheme of human TOP1 with mutants used in panel (D) in red. (D) HEK293 cells were transfected with empty vector (EV) or the His tagged TOP1 WT, Y723F (F) or CPT3KR (3KR) expression plasmid, followed by pre-treatment MG132 then co-treatment with DMSO or 20 μM CPT for 30 min. His-pull down samples were subjected to IB with indicated antibodies. (E) Ub and SUMO proteins are enriched in TOP inhibitors-treated ICE samples. HEK293 cells were exposed to DMSO, 20 μM CPT or 200 μM ETP for 30 min, followed by lysis and CsCl ultracentrifugation using the conditions for ICE assay. Isolated nucleic acids were treated with RNase A, precipitated with ethanol, and analyzed by HPLC-MS/MS. (F) Scheme of the Detection of Ubiquitylated and SUMOylated TOP-DPCs (DUST) assay: 1. Cells are seeded at near confluence and form reversible TOP1ccs (yellow) and TOP2ccs (red); 2. TOP poisons generate TOP-DPCs that are modified by SUMO and Ub PTMs (blue hexagons). Other proteins bound to chromatin are shown as green square; 3. To isolate DPCs, cells are lysed using DNAzol^TM^ and samples ethanol-precipitated, and isolated nucleic acids containing TOP-DPCs are resuspended, sonicated and quantitated; 4. Each sample is either digested with micrococcal nuclease, followed by analysis by SDS electrophoresis and IB with antibodies targeting SUMOs, Ub and TOPs, or subjected to slot-blot and probed with anti-dsDNA antibody as loading control. (G-H) HEK293 cells were exposed to 20 μM CPT or 200 μM ETP for 10, 30, 60, 120, and 240 min, followed by the DUST assay for immunodetection of SUMO-2/3-, SUMO-1-, Ub- and total TOP-DPCs.

To assess whether TOP SUMOylation plays a role in its ubiquitylation, we transfected the TOP1-CPT3KR (K117R, K153R and K103R) construct, which is deficient in SUMOylation due to disruption of its SUMO-consensus motifs (Figure 3C) (Horie et al., 2002; Li et al., 2015). Those mutations not only impaired SUMOylation but also ubiquitylation of TOP1 (Figure 3D), indicating that SUMOylation of TOP1 engages the UPP. The TOP1 catalytic dead (Y723F) mutant (Figure 3C) also underwent SUMO-1 and SUMO-2/3 modifications, albeit to a lesser extent than TOP1 WT (Figure 3D) (Horie et al., 2002). As the Y723F mutant lacking DNA cleavage activity is still capable of binding DNA, this result implies that TOP1 binding is sufficient to elicit its SUMOylation. To demonstrate that TOP1- and TOP2-DPCs are directly conjugated with SUMO-1, SUMO-2/3 and Ub, we performed high performance liquid chromatography and tandem mass spectrum (HPLC-MS/MS) following ICE assay to isolate DNA after treatment of HEK293 cells with CPT or ETP. The MS analysis confirmed that CPT selectively induced TOP1-DPCs and ETP selectively induced TOP2α- and TOP2β-DPCs (Figure 3E), and that SUMO-1, SUMO-2 and Ub were detected in the CPT- and ETP-treated but not vehicle (DMSO)-treated samples.

To directly assess the SUMO and Ub modifications of TOP-DPCs in human cells, we developed an assay for the detection of ubiquitylated and SUMOylated TOP-DPCs (DUST) derived from the RADAR assay (Kiianitsa and Maizels, 2013) (Figure 3F). Briefly, cells treated with TOP poisons (CPT or ETP) are lysed with DNAzol followed by ethanol precipitation. Nucleic acids are collected and digested by micrococcal nuclease. Released TOP-DPCs are resolved by SDS-PAGE electrophoresis and SUMO- and Ub-TOP-DPCs were detected by IB. This DUST assay showed that TOP-DPCs form instantly (within 10 min) after addition of TOP inhibitors (Figure 3G, H, bottom panels), reaching a peak at 30 min, and gradually decreased to become undetectable after 4 hours. SUMO-2/3, SUMO-1 and Ub modifications peaked sequentially (at 10 min, 30 min and 1 h, respectively), indicating that the consecutive SUMO-2/3, SUMO-1 and Ub are conjugated to TOP-DPCs in coordination with each other and with the clearance of TOP-DPCs. In addition, the DUST assays showed that total TOP-DPCs and the PTM-modified TOP-DPCs are induced by TOP inhibitors in a dose-dependent manner (Figure S2A, B).

As both TOP2α and TOP2β are trapped by ETP, the Ub- and SUMO-TOP2-DPCs presumably represent a mixed population comprising both TOP2α- and TOP2β-DPCs. To examine TOP2α and TOP2β separately, we performed DUST assays in WT and TOP2B CRISPR knockout (KO) HeLa cells treated with or without TOP2A siRNA knockdown (KD). Both TOP2α- and TOP2β-DPCs were found to be SUMOylated and ubiquitinylated upon ETP treatment (Figure S2C).

### SUMO and Ub Linkages of TOP1- and TOP2-DPCs in Human Cells

Ubiquitin can be conjugated through one of its 7 lysine residues (K6, K11, K27, K29, K33, K48 and K63) with the K48 or the K11 chains leading to proteasomal degradation (Akutsu et al., 2016). To scrutinize the poly-ubiquitin linkages of TOP-DPCs, we transfected HA-tagged lysine-to-arginine mutants for each ubiquitylation residue and performed DUST assays. K11R-, K48R- and K63R mutations significantly reduced both TOP1- and TOP2-DPC ubiquitylation (Figure S2D, E). The K48 and the K11 linkages are consistent with subsequent proteasomal degradation of TOP-DPCs. Whether the K63 ubiquitylation of TOP-DPCs serves as a platform for the engagement of DDR proteins remains to be determined (Oh et al., 2018).

Given the established role of K11 of SUMO-2/3 in forming polymeric SUMO-2/3 chain (Tatham et al., 2001), we transfected HA-tagged SUMO-2 WT and K11R to determine the TOP-DPC SUMO-2/3 linkages using the DUST assay. The K11R mutation was observed to result in a major decrease in SUMO-2 modification of both TOP1- and TOP2-DPCs (Figure S2F, G, lanes 5, 6).

We also investigated the linkage type of the SUMO-1 chains on TOP-DPCs. Due to lack of internal SUMO consensus motif, it is not well-established whether SUMO-1 by itself is able to form polymers *in vivo*. Because biochemical studies have shown that homogenous SUMO-1 polymer can be assembled through its K7 residue (Yang et al., 2006b), we performed DUST assays in cells transfected with HA-SUMO-1 WT or K7R and found that disruption of the K7 residue did cripple the SUMO-1 polymerization of both TOP1- and TOP2-DPCs (Figure S2F, G, 1, 2).

Acting as an acceptor, SUMO-2/3 moieties can be conjugated with SUMO-1 that terminates SUMO-2/3 chain elongation (Matic et al., 2008). To explore the interplay between SUMO-1 and SUMO-2/3, we downregulated SUMO-2/3 in the SUMO-1-transfected cells and observed shorter forms of SUMO-1-modified TOP-DPCs species (Figure S2F, G, lanes 3). Conversely, KD of SUMO-1 in SUMO-2-transfected cells led to longer forms of SUMO-2-modified TOP-DPCs (Figure S2F, G, lanes 7). These results are consistent with SUMO-2/3 and SUMO-1 sequentially targeting TOP-DPCs with SUMO-2/3 first forming polymeric chains on TOP-DPCs, which are subsequently capped by SUMO-1.

### PIAS4 is a SUMO Ligase for both TOP1- and TOP2-DPCs in Human Cells

To identify the SUMO E3 ligase(s) for TOP1-DPCs, we expressed His-tagged TOP1 in HEK293 cells. Following treatment with CPT or DMSO, we purified TOP1 protein complexes using Ni-NTA to identify co-purifying proteins by LC-MS/MS (Figure 4A, left). Likewise, we expressed FLAG-tagged TOP2α and TOP2β in cells treated with ETP or DMSO, performed FLAG-IP and analyzed TOP2-containing protein complexes by MS (Figure 4A, right). Strikingly, all samples were consistently enriched with PIAS4 (Figure 3A, bottom), a member of Siz/PIAS SUMO ligase family, which SUMOylates DDR proteins including RPA and MDC1 for DSB repair (Galanty et al., 2009). To validate this result, we reciprocally pulled down PIAS4-containing protein complexes by expressing FLAG-PIAS4 in HEK293 cells. MS retrieved TOP1, TOP2α and TOP2β (Figure 4B). IP-IB and Proximity Ligation Assay (PLA) also showed that PIAS4 interacts with TOP1 and TOP2 in the presence or absence of TOP inhibitors (Figure S3A-E).

**Figure 4.**
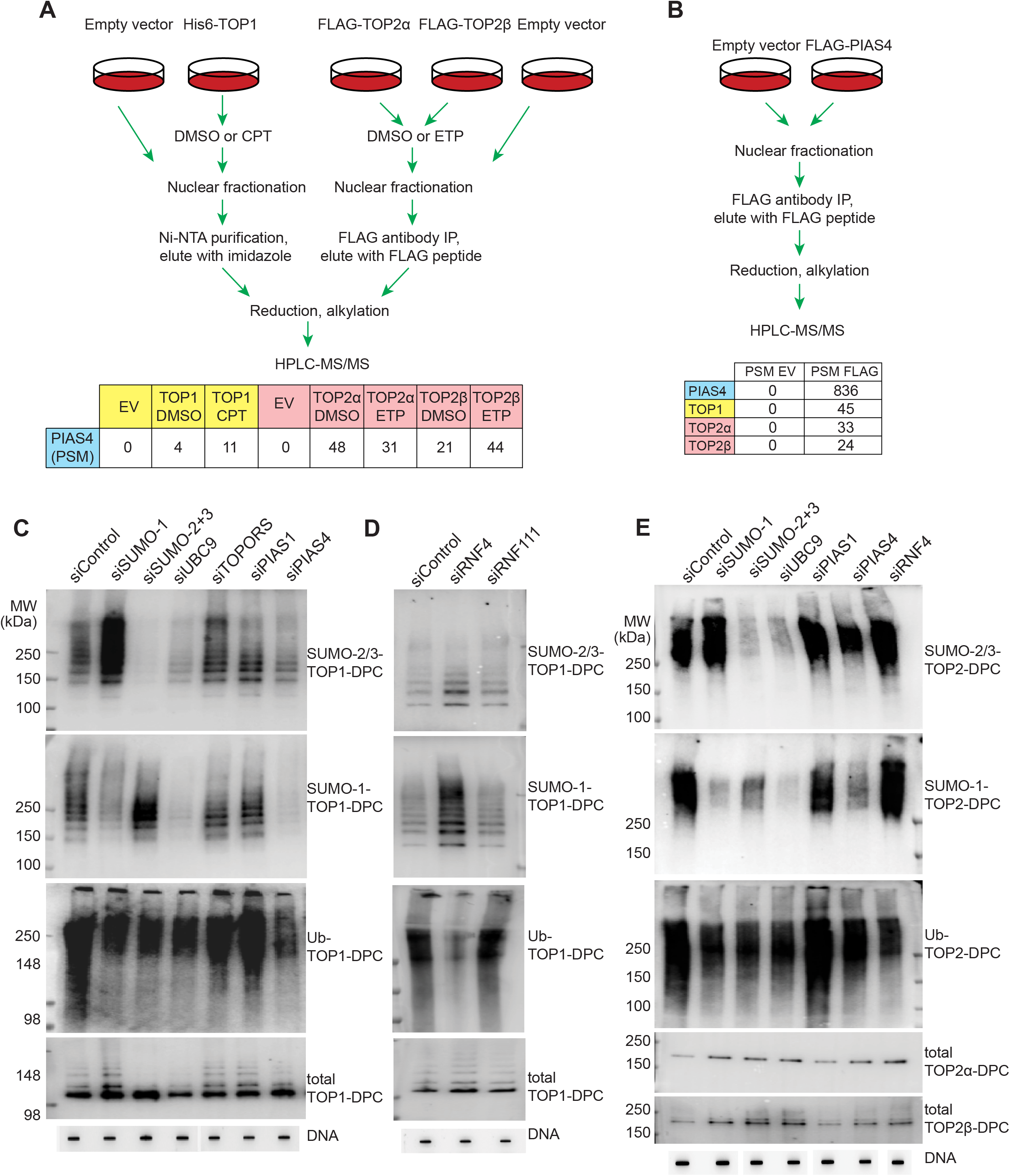
Identification of PIAS4 as SUMO-ligase and RNF4 as STUbL for TOP1- and TOP2-DPCs in Human Cells. (A) Schemes of TOP1-His-IP pull-down, TOP2α- and TOP2β-FLAG-IP pull-down for LC-MS/MS. (B) Scheme of PIAS4 FLAG-IP pull-down for LC-MS/MS. (C) HEK293 cells were transfected with indicated siRNAs, followed by treatment with 20 μM CPT for 30 min (same conditions used for the following DUST assays unless otherwise indicated). Representative DUST assay for immunodetection of SUMO-2/3-, SUMO-1-, Ub- and total TOP1-DPCs. (D) HEK293 cells were transfected with control, RNF4 or RNF111 siRNA, followed by CPT treatment and DUST assay for detection of SUMO-, Ub- and total TOP1-DPCs. (E) HEK293 cells were transfected with indicated siRNAs followed by treatment with 200 μM ETP for 30 min (same conditions used for the following DUST assays unless otherwise indicated). DUST assay for immunodetection of SUMO-2/3-, SUMO-1-, Ub- and total TOP2-DPCs.

PIAS4 downregulation (Figure S3D) decreased the levels of both SUMO-1- and SUMO-2/3-conjugated TOP-DPCs whereas KD of PIAS1, another PIAS family SUMO ligase did not alter SUMOylation of TOP-DPCs in the DUST assays (Figure 4C, E).

To establish that PIAS4 catalyzes TOP SUMOylation, we performed biochemical assays in reactions containing SUMO-1 or SUMO-2, SUMO E1, E2, PIAS4 and recombinant topoisomerases. By probing with anti-SUMO-1 and SUMO-2/3 antibodies, we established that PIAS4 can catalyze TOP1 with preference for SUMO-1 conjugation over SUMO-2 conjugation (Figure S4A). Analogously, PIAS4 also catalyzed SUMO-1 and SUMO2 conjugation of TOP2α and TOP2β (Figure S4B, C)

### RNF4 is a STUbL for both TOP1- and TOP2-DPCs in Human Cells

Because PIAS4 KD was also observed to attenuate TOP-DPC ubiquitylation (Figure 4C, E), we surmised that SUMOylation of TOP-DPC is a trigger for their ubiquitylation. Thus, we tested RNF4 as the human ortholog of Slx5-Slx8 (see above and Figure 1) for TOP-DPC ubiquitylation in human cells. DUST assays showed that RNF4 KD reduced TOP-DPC ubiquitylation while enhancing TOP-DPC SUMOylation (Figure 4D, E). Silencing of RNF111, the other reported mammalian STUbL (Poulsen et al., 2013) did not impact TOP1-DPCs ubiquitylation (Figure 4D).

To validate the activity of RNF4 biochemically, we performed Ub conjugation assays with recombinant topoisomerases, Ub, Ub E1, RNF4 as well as E2 enzyme UbcH5α, which cooperates with RNF4 for ubiquitylation (Plechanovova et al., 2012). IB with anti-Ub antibody showed that TOP ubiquitylation increased in a RNF4-dependent manner (Figure S4D-F). To establish that RNF4 targets SUMOylated topoisomerases, we performed experiments coupling *in vitro* SUMOylation and ubiquitylation. RNF4 displayed much stronger ubiquitylating activity toward SUMO-modified TOPs than toward unmodified TOPs (Figure S4D-F).

### Coordination of PIAS4 and RNF4 for SUMOylation and Ubiquitylation of TOP-DPCs

To explore how PIAS4 and RNF4 coordinate the SUMO-Ub pathway, we overexpressed PIAS4 and RNF4 in HCT116 cells. DUST assays showed that PIAS4 transfection stimulated both TOP-DPC SUMOylation and ubiquitylation, and that RNF4 transfection largely potentiated ubiquitylation of TOP-DPCs (Figure 5A, B, lanes 2, 3). Expression of PIAS4 in PIAS4 CRIPSR KO HCT116 cells in part restored SUMOylation and ubiquitylation whereas ectopic expression of RNF4 in PIAS4 KO cells failed to induce TOP-DPC ubiquitylation (Figure 5A, B, lanes 5, 6; Figure S5A). We conclude that RNF4 ubiquitylates TOP-DPCs in a PIAS4-dependent manner.

**Figure 5.**
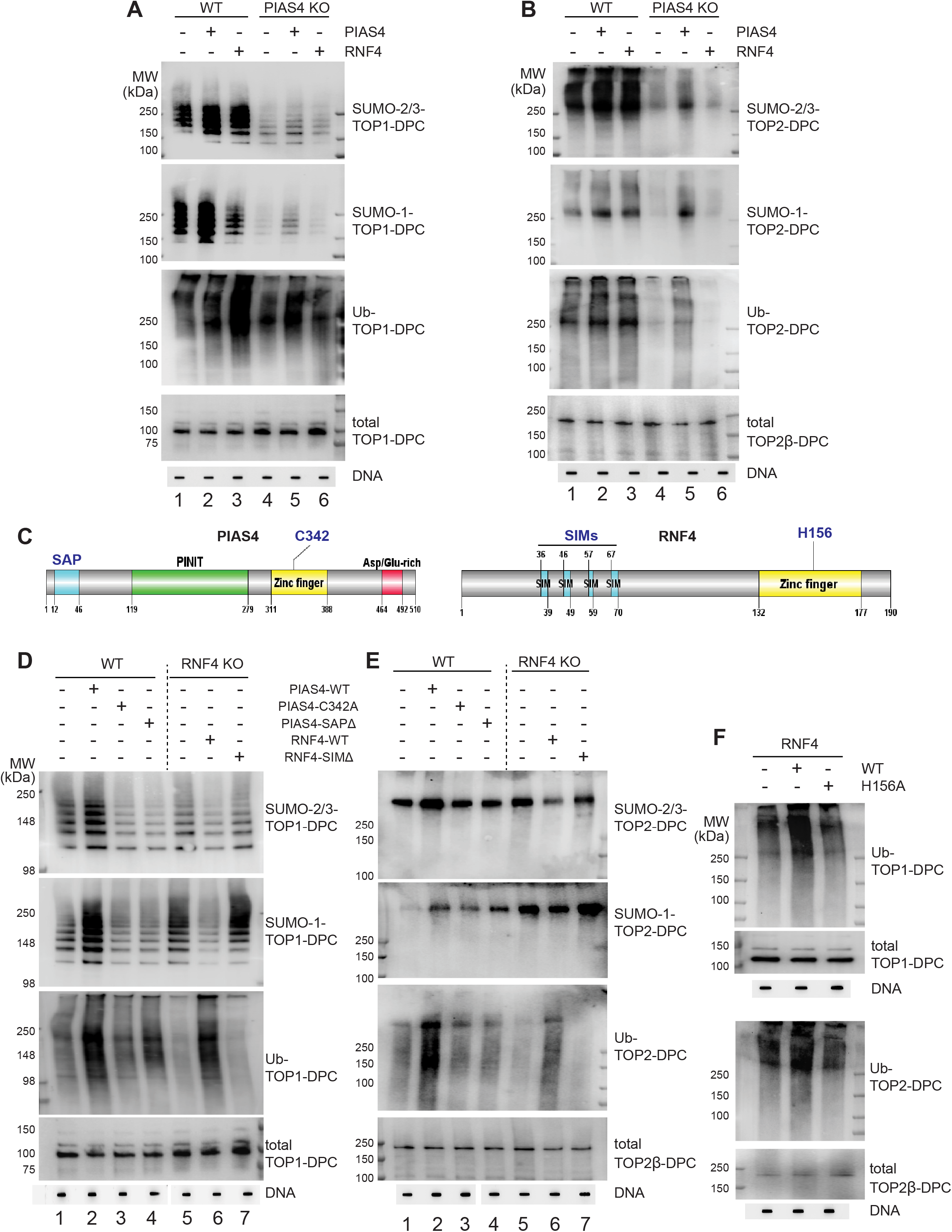
Active Motifs of PIAS4 and RNF4 are Required for the SUMOylation and Ubiquitylation of TOP-DPCs. (A) HCT116 WT and PIAS4-KO cells transfected with indicated plasmids were treated with CPT and analyzed by DUST assay for detection of SUMO-2/3-, SUMO-1- and Ub-TOP1-DPCs. (B) Same as panel (A) except that cells were treated with ETP to induce TOP2-DPCs. (C) Left, Scheme of human PIAS4. Right, Scheme of human RNF4. (D) MCF7 WT and RNF4 KO cells were transfected with indicated plasmids, followed by CPT treatment and analyzed by DUST assay for detection of SUMO-2/3-, SUMO-1- and Ub-TOP1-DPCs. (E) Same as panel (D) except that cells were treated with ETP to induce TOP2-DPCs. (F) Upper panel, MCF7 RNF4-KO cells were transfected with RN4 WT or H156A plasmids, followed by CPT treatment (20 μM, 1 h) and DUST assay for immunodetection of Ub-TOP1-DPCs. Lower panel: Same as upper panel except that ETP (200 μM, 1 h) was used to induce TOP2-DPCs.

PIAS4 has been shown to localize to DSB sites by binding DNA via its N-terminal SAP domain (Figure 5C, left) in association with DDR proteins (Galanty et al., 2009). Accordingly, PIAS4 SAPΔ as well as the PIAS4 catalytic-dead (C342A) transfected MCF7 cells were defective in TOP-DPC SUMOylation and ubiquitylation (Figure 5D, E, lanes 3, 4, Figure S5C), indicating that SUMOylation of TOP-DPCs is dependent on the recruitment of PIAS4 to chromatin.

To test whether RNF4 ubiquitylates TOP-DPCs through its SUMO interacting motifs (SIMs) (Figure 5C, right) (Kung et al., 2014), we transfected RNF4 WT and SIMΔ plasmids in RNF4 KO MCF7 cells. The SIM mutant failed to induce TOP-DPC ubiquitylation in RNF4 KO cells as opposed to the RNF4 WT (Figure 5D, E, lanes 7; Figure S5B, D). Similarly, a RNF4 catalytic-dead mutant (H156A) was unable to induce TOP-DPC ubiquitylation in RNF4 KO cells (Figure 5F, Figure S5D). These results were corroborated by PLA assays showing that PIAS4 SAPΔ and RNF4 SIMΔ fail to colocalize with TOP1 and TOP2α (Figure S5E-F). We conclude that PIAS4 is recruited to TOP-DPCs through its DNA binding domain and TOP-DPC SUMOylation recruits RNF4 through its SIM motifs, leading to TOP-DPC ubiquitylation.

### Connecting SUMOylation and SUMO-dependent Ubiquitylation of TOP-DPCs with DNA transactions and DDR in Human Cells

To investigate whether DNA transactions and the DNA damage response (DDR kinases) play a role in the SUMOylation and ubiquitylation of TOP-DPCs, we performed DUST assays in HEK293 cells treated with the DNA-PKcs inhibitor VX984, the ATM inhibitor KU55399, the ATR inhibitor AZD6738, the replication inhibitor aphidicolin (APH) or the transcription inhibitor DRB prior to CPT or ETP treatment. None of the inhibitors affected the SUMOylation of TOP-DPCs (Figure 6A, B), indicating that TOP-DPC SUMOylation is induced prior to collision between TOP-DPCs and DNA transaction machineries and activation of DDR. These findings may explain why SUMO modifications of TOPs transpire within minutes of the DPC formation.

**Figure 6.**
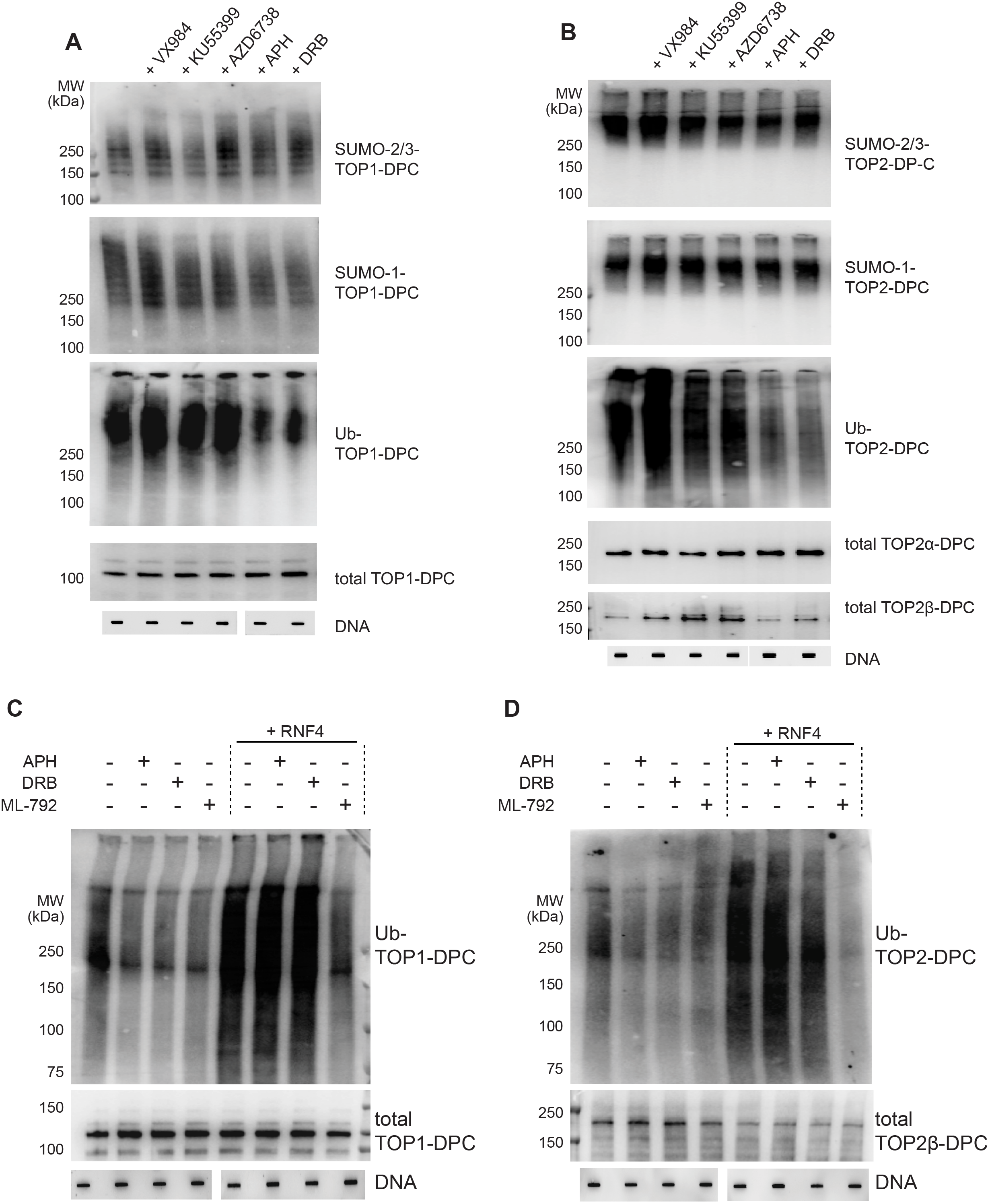
SUMOylation and SUMO-dependent ubiquitylation of TOP-DPCs and DNA Transactions and DDR in Human Cells. (A) HEK293 cells were pre-treated with the DNA-PK inhibitor VX984 (10 μM, 6 h), the ATM inhibitor KU55399 (10 μM, 6 h), the ATR inhibitor AZD6738 (10 μM, 6 h), the replication inhibitor aphidicolin (APH, 10 μM, 2 h), the transcription inhibitor DRB (100 μM, 2 h) before co-treatment with CPT. Cells were then subjected to DUST assay for detection of SUMO-2/3-, SUMO-1-, Ub-, and total TOP1-DPCs. (B) Same as panel (A) except that ETP was used to induce TOP2-DPCs. (C) HEK293 cells were transfected with or without RNF4 plasmid as indicated. Following pre-treatment with APH, DRB or ML-792, 20 μM CPT was added for an additional 1 h. DUST assay was used to detect Ub-TOP1-DPCs. (D) Same as panel (C) except that ETP (200 μM, 1h) was used to induce TOP2-DPCs.

While TOP-DPC SUMOylation was insensitive to ATM, ATR and DNA-PK inhibitors, ubiquitylation was reduced by replication or transcription inhibition (APH and DRB, Figure 6A, B). However, APH and DRB did not exhibit any significant impact on TOP-DPC ubiquitylation potentiated by RNF4 overexpression (Figure 6C, D), implying that there are other ubiquitylation pathways activated upon collision of replication and transcription with TOP-DPCs (Desai et al., 2003; Li et al., 2003; Lin et al., 2009; Xiao et al., 2003).

### RNF4 Drives the Proteasomal Degradation of TOP-DPCs in Human Cells

To link RNF4-mediated ubiquitylation to TOP-DPC proteasomal degradation, we conducted ICE assay in MCF WT cells and found that overexpression of RNF4 reduced TOP-DPCs (Figure 7A-C; Figure S7A, B). This reduction was suppressed by MG132, indicating that RNF4 induces TOP-DPC removal through the proteasome. In addition, MG132 failed to further increase TOP-DPCs in MCF RNF4 KO cells, suggesting an epistatic relationship between RNF4 and the proteasome in TOP-DPC removal (Figure 7A-C; Figure S6A, B).

**Figure 7.**
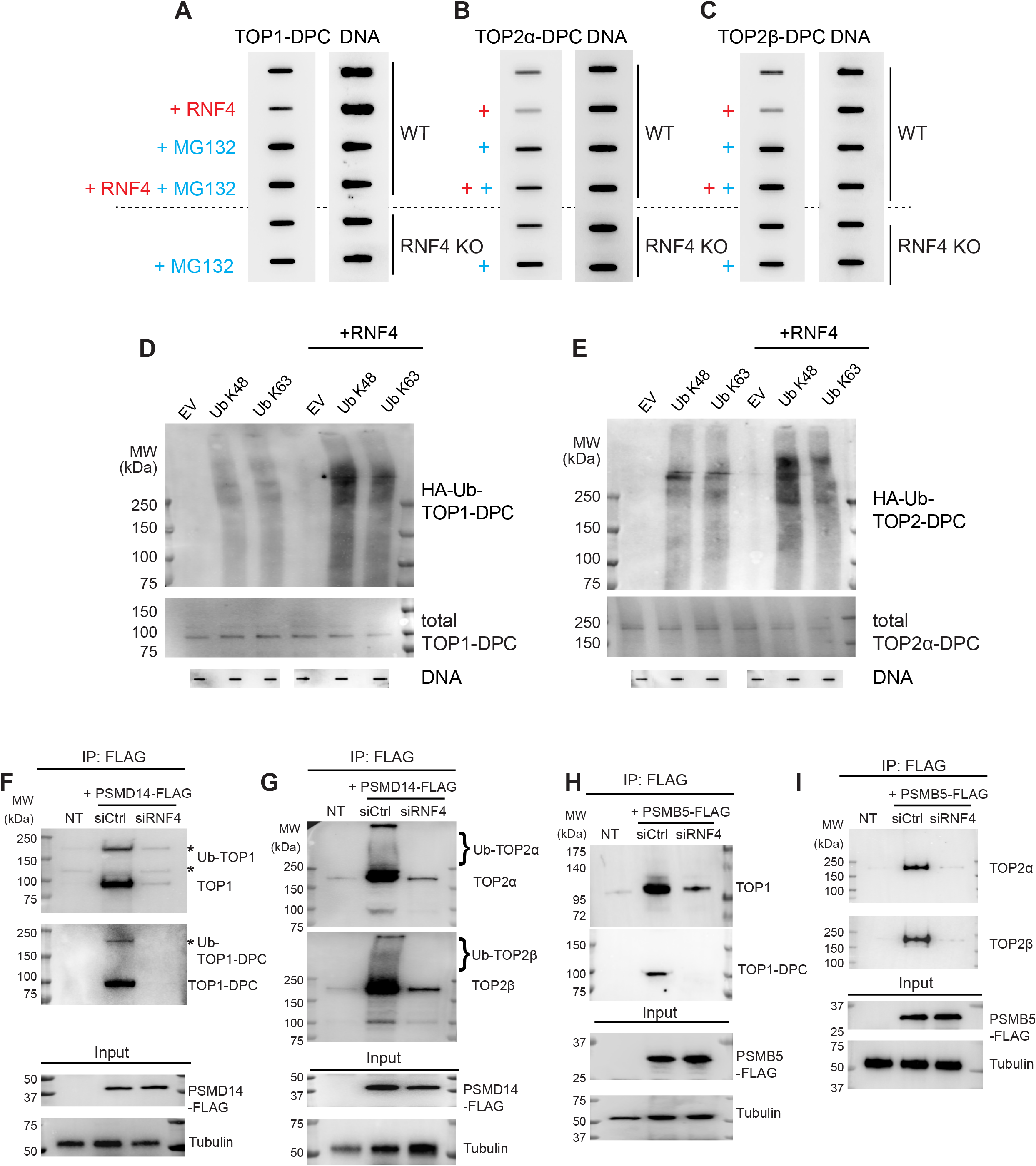
RNF4 Drives TOP-DPC Proteasomal Degradation. (A) MCF7 WT cells were transfected with or without RNF4 and pre-treated with or without MG132, followed by CPT (1 μM, 2 h) co-treatment. RNF4-KO cells were pre-treated with or without MG132, then co-treated with CPT (1 μM, 2 h) followed by ICE assay for TOP1-DPC detection. (B) Same as panel (A) except that CPT was replaced by ETP (10 μM, 2 h) for ICE assay detecting TOP2α-DPCs. (C) Same as panel (B) except that IB was performed with TOP2β antibody. (D) HEK293 cells were transfected with empty vector, HA-Ub K48 or HA-K63, followed by transfection with RNF4 as indicated. Following CPT treatment (20 μM, 1 h), cells were subjected to DUST assay using anti-HA antibody. (E) Same as panel (D) except that cells were treated with ETP (200 μM, 1h) for HA-Ub-TOP2-DPC detection. (F) HEK293 cells transfected with control siRNA or RNF4 siRNA and PSMD14-FLAG expression plasmid were pre-treated with MG132 before co-treatment with CPT (20 μM, 1 h). Cells were then subjected to FLAG-IP and IB with anti-TOP1 and -TOP1-DPC antibodies. NT, no transfection. (G) Same as panel (F) except that cells were treated with ETP (200 μM, 1h) and that IB was performed with anti-TOP2α and β antibodies. (H) HEK293 cells transfected with control siRNA or RNF4 siRNA and PSMB5-FLAG expression plasmid were pre-treated with MG132 before co-treatment with CPT (20 μM, 1 h). Cells were then subjected to FLAG-IP and IB with anti-TOP1 and -TOP1-DPC antibodies. (I) Same as panel (H) except that cells were treated with ETP (200 μM, 1h and that IB was performed with anti-TOP2α and β antibodies.

As we determined that TOP-DPCs are mainly modified by Ub at K48 and 63, we tested which Ub linkage(s) is mediated by RNF4. To do so, we transfected HEK293 cells with single lysine Ub constructs (the other lysines are converted into arginine) HA-Ub K48 or HA-Ub K63. Following RNF4 transfection and DUST assays, we found that RNF4 significantly facilitated K48 ubiquitylation but not K63 ubiquitylation of TOP-DPCs (Figure 7D, E). This result suggests that RNF4 primes TOP-DPCs for proteasomal degradation by catalyzing the K48 poly-ubiquitylation.

Next, we tested potential direct interactions of the proteasome with TOP-DPCs. During degradation, the proteasome non-ATPase regulatory subunit 14 (PSMD14) deubiquitylates substrates to enable their access into the proteasome catalytic core (Bard et al., 2018). To determine whether PSMD14 interacts with TOP-DPCs, we transfected a PSMD14-FLAG expression plasmid into HEK293 cells and performed IP after MG132 and CPT or ETP co-treatment. We observed multiple bands that have higher molecular weights than the actual TOP1 by IB using anti-TOP1 and -TOP1-DPC antibodies in CPT-treated IP samples (Figure 7F). Similarly, smeared bands above TOP2 were detected in ETP-treated IP samples using anti-TOP2 antibodies (Figure 7G). As the proteasome lacks deSUMOylating activity, it is conceivable that SUMO chains conjugated to TOPs are removed before proteasome processing. We conclude that the observed upper shifted or smeared bands of TOPs indicate poly-Ub-TOP species. Downregulation of RNF4 largely diminished the interaction (Figure 7F, G), suggesting that PSMD14 binds RNF4-generated Ub-TOP for deubiquitylation, allowing the entry of TOPs into the proteasome core particle.

We next transfected HEK293 cells with a plasmid overexpressing the 20S proteasome core subunit β type-5 (PSMB5) possessing chymotrypsin-like activity. Following MG132 and CPT or ETP co-treatment, FLAG-IP experiments showed that PSMB5 only interacted with unmodified TOPs (Figure 7H, I), suggesting that TOPs are translocated to the catalytic core following their deubiquitylation. KD of RNF4 attenuated the interaction between PSMB5 and TOPs (Figure 7H, I), substantiating the role of RNF4 in driving the proteasomal degradation of TOP-DPCs. By probing the CPT-treated IP samples with antibody targeting TOP1-DPC, we confirmed that PSMB5 interacts TOP-1-DPC in a RNF4-dependend manner, directly suggesting that the 20S core complex targets covalent DNA-bound TOPs for proteolysis. As proteasome inhibition precludes γH2AX induction by TOP-DPCs presumably by perpetuating non-proteolyzed TOP-DPCs that conceal the TOP-linked DSBs, we examined whether PIAS4 and RNF4 are required for γH2AX. IF in U2OS cells treated with CPT or ETP revealed that PIAS4- and RNF4-KD reduced the levels of γH2AX (Figure S6C-F), suggesting that PIAS4-RNF4-mediated proteasomal degradation of TOP-DPCs enables activation of DDRs such as γH2AX.

## DISCUSSION

Our study presents an evolutionally conserved SUMO-Ub-proteasome pathway of both TOP1 and TOP2-DPCs for yeast and human cells. This pathway consists of a choreographed SUMO-Ub-proteasome axis driven by human PIAS4 and yeast Siz1 as SUMO ligases and human RNF4 and yeast Slx5/Slx8 as STUbLs. For these studies, we designed a novel assay (DUST) to detect Ub- and SUMO-TOP-DPCs.

We show that SUMO-2/3, SUMO-1 and ubiquitin are conjugated to TOP-DPCs sequentially in human cells. SUMO-2/3 conjugation occurs first, with a maximum in less than 10 minutes, while SUMO-1 and Ub peak later (at 30 and 60 minutes, respectively). Our finding that SUMO-2/3 downregulation prevents both SUMO-1 and Ub modifications of TOP-DPCs is consistent with rate-limiting role for SUMO-2/3 modifications in TOP-mediated damage signaling and repair. SUMO-1 KD resulted in increased SUMO-2/3-TOP-DPC species, denoting a role of SUMO-1 in limiting SUMO-2/3 polymer elongation and exemplifying how mammalian cells differentially employ two types of SUMO modifications. This division of labor is relevant to a recent study showing that SUMO-2/3 but not SUMO-1 modification of human TOP2 enables TDP2 to excise TOP2-DPCs without proteolyzing the bulk of the protein (Schellenberg et al., 2017). Our data also show that the expeditious SUMOylation of TOP-DPCs is not coupled with replication or transcription, and that excessive TOP1 binding to chromatin without DNA cleavage is sufficient to evoke TOP1 SUMOylation. The extensive involvement and regulatory role of SUMOylation in DDR extends beyond TOP-DPCs. Indeed, TOP-DPC SUMOylation is likely part of a more extensive SUMO response surrounding DNA damage sites (Jackson and Durocher, 2013) and impacting other DDR and repair proteins including TDP1 (Hudson et al., 2012), 53BP1, BRCA1 and Rad52 (Galanty et al., 2009; Jackson and Durocher, 2013; Morris et al., 2009).

Our study reveals PIAS4 as SUMO ligase for human TOP-DPCs in addition to its established roles for DDR protein SUMOylation. We also show that in yeast TOP-DPC SUMOylation is carried out by Siz1, the ortholog of PIAS4. Genetic suppression of PIAS4 in human cells as well as biochemical assays demonstrate the activity of PIAS4 as E3 ligase toward TOP1, TOP2α and TOP2β. Notably, our work also reveals that PIAS4 and TOPs interact even in the absence of TOP inhibitors, implying that PIAS4 colocalizes with topoisomerases in chromatin and carries out the prompt SUMOylation of TOP-DPCs once they become abortive. This possibility is supported by our finding that the DNA binding domain (SAP) of PIAS4 is critical for TOP-DPC SUMOylation. Thus, we propose that the DNA binding of PIAS4 next to the TOP-DPC activates PIAS4 and TOP-DPC SUMOylation. Together, these experiments demonstrate the importance of PIAS4 (and Siz1) in the early response to TOP-DPCs.

Our study connects SUMOylation of TOP-DPCs to their ubiquitylation through the STUbL RNF4 in humans and Slx5-Slx8 in yeast. Suppressing SUMOylation genetically and pharmacologically reduced TOP-DPC ubiquitylation. RNF4/Slx5-Slx8 are pivotal components in this SUMO-Ub axis, as demonstrated by our finding that depletion of RNF4 in human cells and Slx5-Slx8 in yeast reduce the levels of Ub-TOP-DPCs and elevate SUMO-TOP-DPCs. As a well-studied STUbL, RNF4 is activated upon formation of poly-SUMOylated substrates through its SIM domains and to contain a DNA binding motif within its RING domain (Groocock et al., 2014; Rojas-Fernandez et al., 2014). Of note, our DUST experiments show that PIAS4-mediated SUMOylation is activated independently of replication, transcription and DDRs, and that RNF4-catalyzed TOP-DPC ubiquitylation only requires SUMOylation. These data signify a role of SUMOylation as the first line of defense of toxic TOP-DPCs by priming them for RNF4-mediated proteasomal degradation by. A potentially important determinant of PIAS4 and RNF4 activation could be their binding to exposed DNA segments next to TOP-DPCs. RNF4 appears to be a key component of the DDR impacting both histones and DNA repair proteins including BRCA1, MDC1, FANCD2, and 53BP1 (Gibbs-Seymour et al., 2015; Groocock et al., 2014; Kumar et al., 2017; Poulsen et al., 2013; Vyas et al., 2013), suggesting that the ubiquitylation of TOP-DPCs is part of a larger DDR response at the DNA damage sites.

The importance of the proteasome in degrading TOP-DPCs has been established by many studies prior to this one [reviewed in (Pommier et al., 2016)]. Yet, our study provides novel molecular insights in this process by demonstrating the cellular binding of TOP1, TOP2α and TOP2β to at least two components of the 26S proteasome: PSMD14 for deubiquitylation and PSMB5 for proteolytic digestion. We also show that the RNF4-mediated Ub linkage to TOPs is through lysine 48 chain, which is required for of the interaction between proteasome subunits and TOP-DPCs. Further studies are warranted to establish whether the SUMO-Ub-proteasome pathway described here extends to other enzymatic DPCs such as those generated by DNA methyltransferases and chemically by formaldehyde or chemotherapeutic crosslinking agents such platinum derivatives. (Borgermann et al., 2019; Larsen et al., 2019; Sparks et al., 2019).

## SUPPLEMENTAL FIGURE LEGENDS

**Figure S1.**
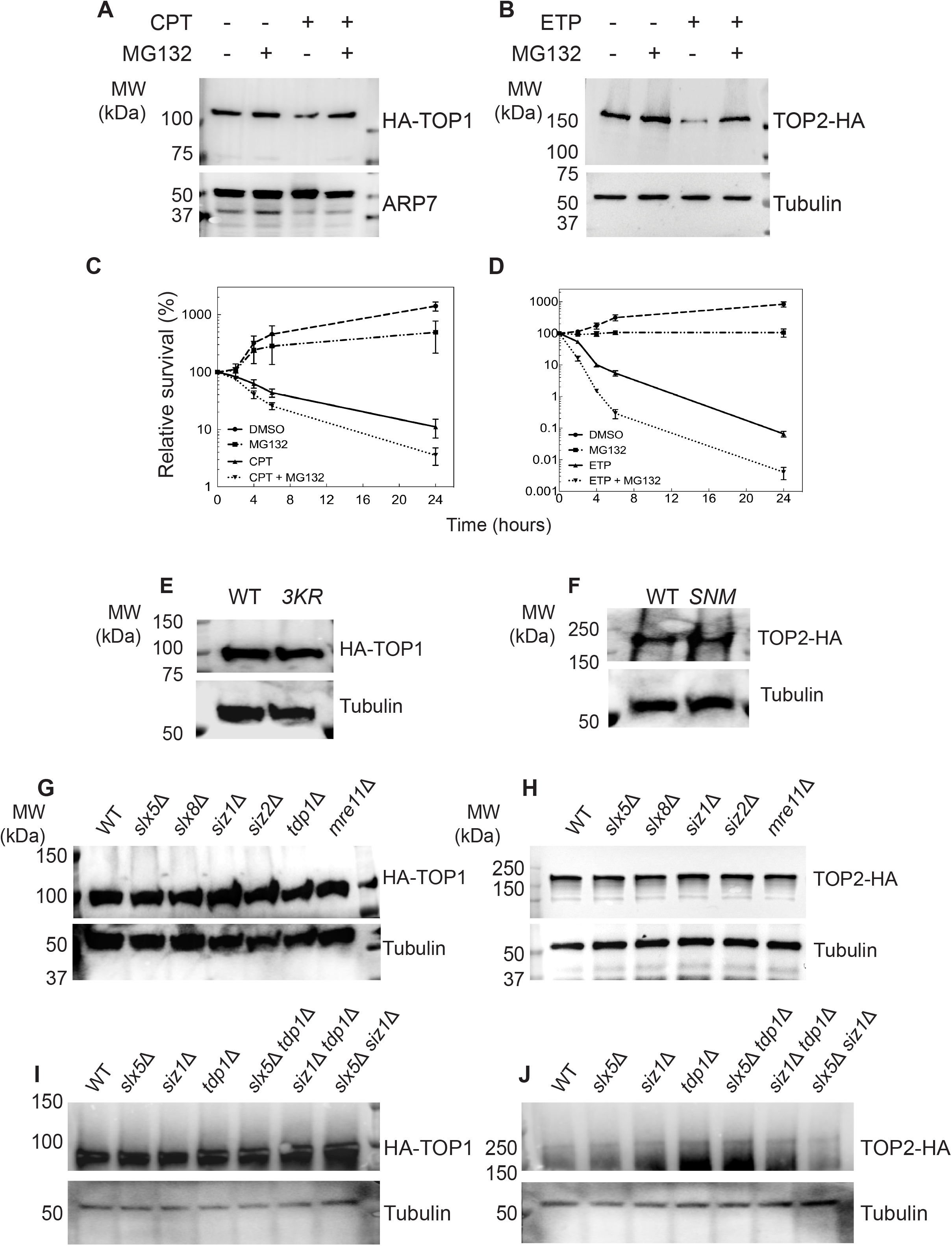
The Proteasome Participates in Repair of TOP-mediated DNA Damage in Yeast. (A) TOP1 levels in YMM10 cells carrying pYX112 HA-TOP1 plasmid after treatments with DMSO, 10 μM MG132, 10 μg/ml CPT or CPT + MG132 for a total of 4 h. Lysates were analyzed by WB with anti-HA and anti-ARP7 antibodies. (B) TOP2 levels in YMM10 cells carrying pDED1 TOP2-3×HA plasmid following treatments with DMSO, MG132, 100 μg/ml ETP or ETP + MG132 for a total of 4 h. Lysates were analyzed by WB with anti-HA and anti-α tubulin antibodies. (C) YMM10 cells overexpressing HA-TOP1 were exposed to DMSO, 10 μM MG132, 5 μg/ml CPT or CPT + MG132 at 30°C for different times (2, 4, 6, 24 h). Aliquots were removed, diluted, and plated to SD-ura plates for clonogenic assays. Cell survival is expressed as the percentage of surviving cells at the time points relative to the viable titer at the time drug was added (t = 0). Error bars represent the standard deviation of three independent experiments. Data points without error bars have error bars that are smaller than the symbol drawn on the graph. (D) YMM10 cells overexpressing TOP2-HA were exposed to DMSO, 10 μM MG132, 50 μg/ml ETP or ETP + MG132 at 30°C for different times (2, 4, 6, 24 h) for clonogenic assay as described in panel (C). (E) WB showing TOP1 levels in YMM10 strains carrying the HA-TOP1 WT or 3KR plasmid. (F) WB showing TOP2 levels in YMM10 strains carrying the TOP2-3×HA WT or *SNM* plasmid. (G) WB showing TOP1 levels in indicated BY4741 strains carrying the HA-TOP1 plasmid. (H) WB showing TOP2 levels in indicated BY4741 strains carrying the TOP2-3×HA plasmid. (I) WB showing TOP1 levels in indicated BY4741 strains carrying the HA-TOP1 plasmid. (J) WB showing TOP2 levels in indicated BY4741 strains carrying the TOP2-3×HA plasmid.

**Figure S2.**
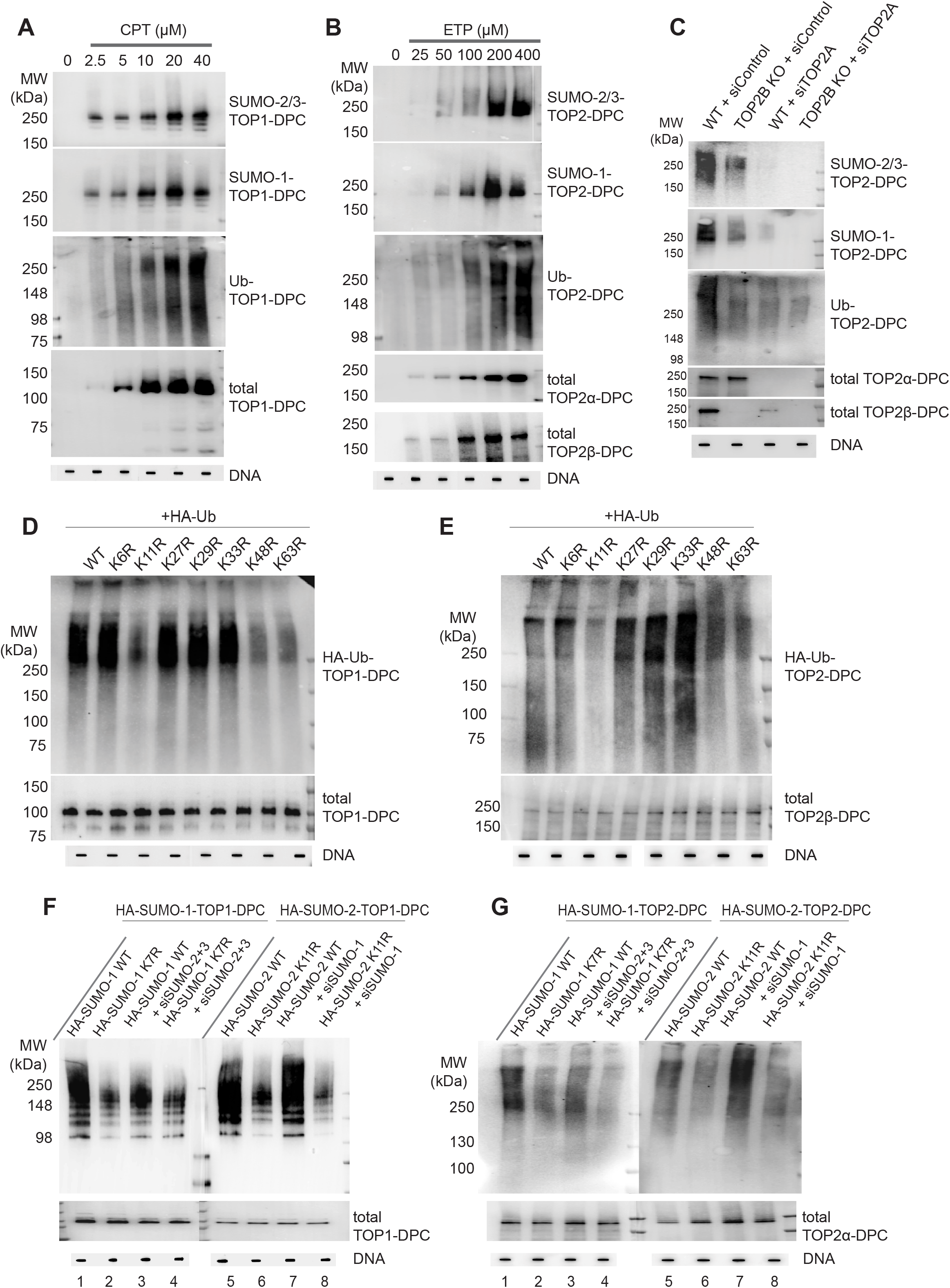
SUMO and Ubiquitin Linkages of TOP1- and TOP2-DPCs in Human Cells. (A) HEK293 cells were exposed to increasing concentration of CPT for 30 min, followed by DUST assay for immunodetection of SUMO-2/3 modified TOP1-DPCs, SUMO-1 modified TOP1-DPCs, ubiquitylated TOP1-DPCs and total TOP1-DPCs. (B) HEK293 cells were exposed to ETP at a series of concentrations for 30 min, followed by DUST assay for immunodetection of SUMO-2/3 modified TOP2-DPCs, SUMO-1 modified TOP2-DPCs, ubiquitylated TOP2-DPCs and total TOP2-DPCs. (C) HeLa WT and TOP2B KO cells transfected with control siRNA or TOP2A siRNA were treated with ETP (200 μM, 30 min) before performing DUST assay for detection of SUMO-1 modified TOP2-DPCs, SUMO-2/3 modified TOP2-DPCs, ubiquitylated TOP2-DPCs and total TOP2-DPCs. (D) HEK293 cells were transfected with the indicated plasmids to overexpress HA-tagged WT and mutant ubiquitin proteins, followed by CPT treatment (20 μM, 1h) and DUST assay for ubiquitylated TOP1-DPC detection. (E) HEK293 cells were transfected with various plasmids to overexpress HA-tagged WT and mutant ubiquitin proteins, followed by ETP treatment (200 μM, 1h) and DUST assay for ubiquitylated TOP2-DPC detection using anti-HA antibody. (F) HEK293 cells were transfected with indicated plasmids and siRNAs, followed by CPT treatment (20 μM, 1h) and DUST assay for SUMO-1 and SUMO-2 modified TOP1-DPC detection using HA antibody. (G) HEK293 cells were transfected with indicated plasmids and siRNAs, followed by ETP treatment (200 μM, 1h) and DUST assay for SUMO-1 and SUMO-2 modified TOP2-DPC detection using anti-HA antibody.

**Figure S3.**
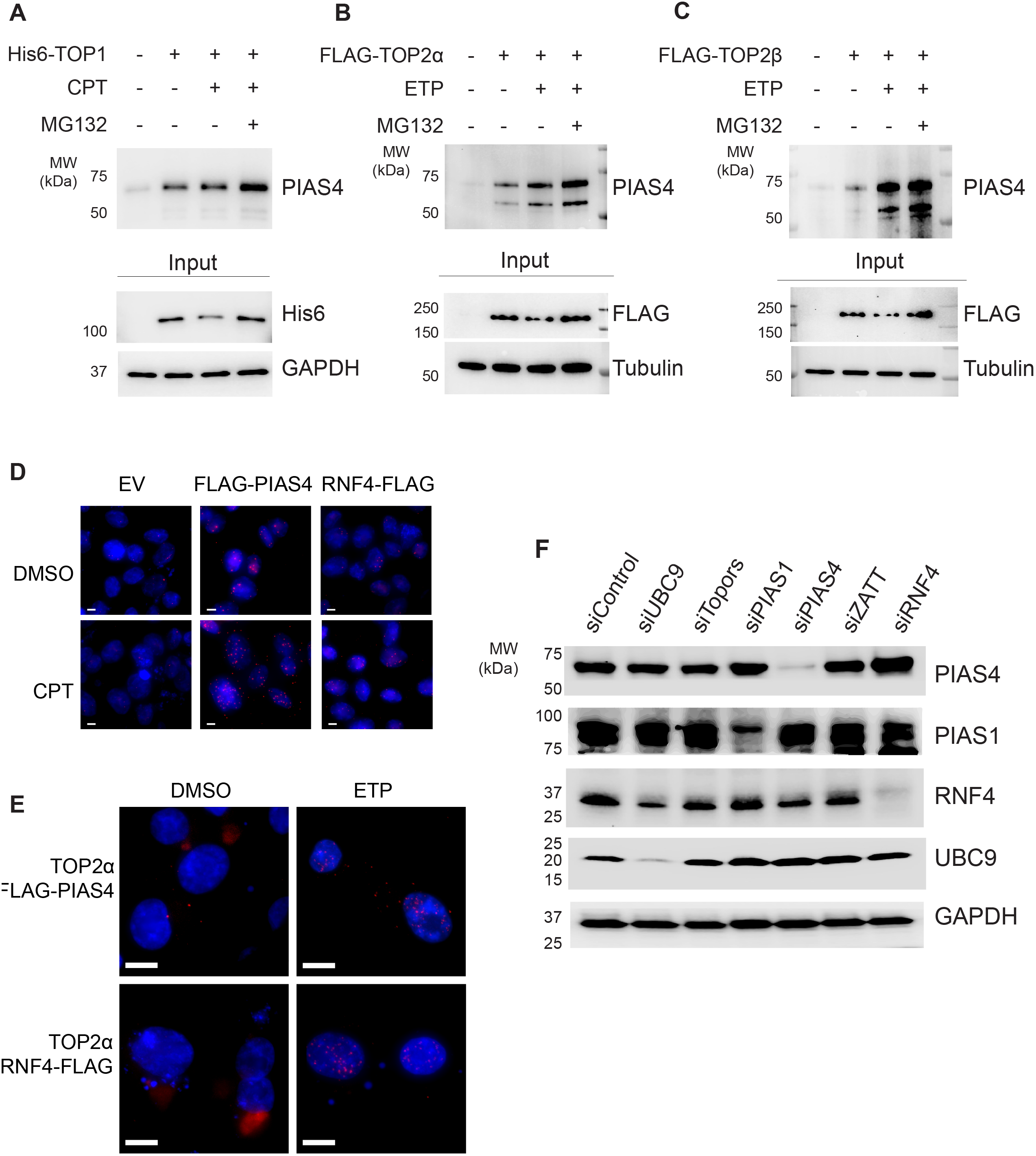
PIAS4 Physically Interacts with TOP1, TOP2◻ and TOP2◻ in the Absence or Presence of TOP Inhibitors. (A) Cells were transfected with the 6×His tagged TOP1 expression splasmid, followed by treatments with DMSO or 20 μM CPT in absence or presence of 10 μM MG132 for 30 min. Native His-pull down was performed as described under “Materials and Methods.” His-pull down samples and cell lysates (input) were then subjected to IB using indicated antibodies. (B) Cells were transfected with the FLAG tagged TOP2α expression plasmid, followed by treatments with DMSO or 200 μM ETP in absence or presence of MG132 for 30 min. Native FLAG-IP was performed as described under “Materials and Methods.” IP samples and cell lysates (input) were then subjected to IB using indicated antibodies. (C) Cells were transfected with the FLAG tagged TOP2β expression plasmid, followed by treatments with DMSO or ETP in the absence or presence of MG132 for 30 min. IP samples and cell lysates were subjected to IB using indicated antibodies. (D) U2OS cells were transfected with empty vector (EV), FLAG-PIAS4 or RNF4-FLAG expression plasmids then subjected to DMSO or CPT treatment (20 μM, 30 min), followed by proximity ligation assay using anti-TOP1 antibody and anti-FLAG antibody. Scale bar represents 10 microns. (E) U2OS cells were transfected with FLAG-PIAS4 or RNF4-FLAG expression plasmids then subjected to DMSO or ETP treatment (200 μM, 30 min), followed by PLA using anti-TOP2α antibody and anti-FLAG antibody. Scale bar represents 10 microns. (F) Cells were transfected with indicated siRNAs, followed by WB of the whole cellular lysates to validate downregulation of each protein.

**Figure S4.**
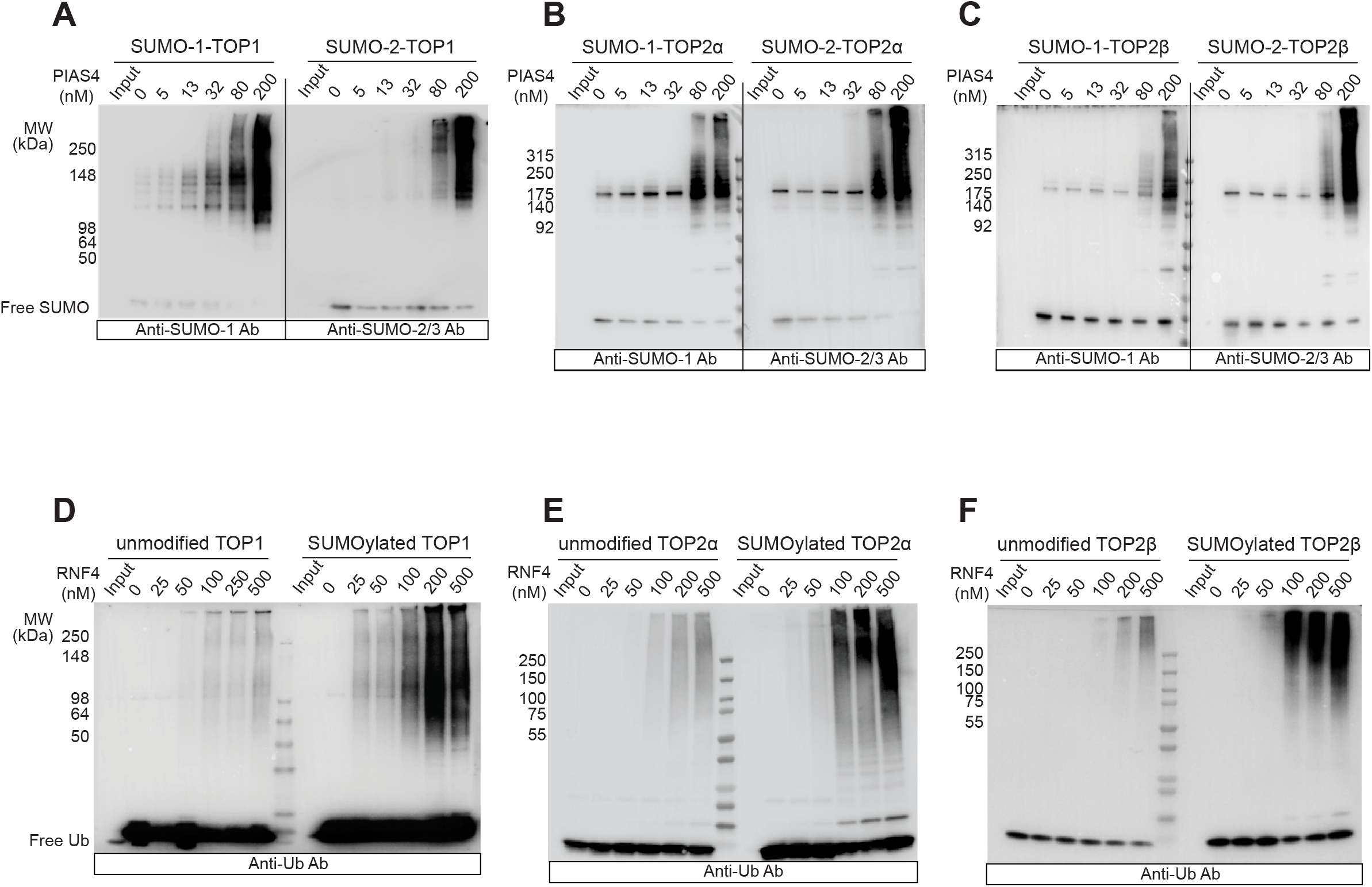
in vitro SUMOylation and Ubiquitylation of Human Topoisomerases by PIAS4 and RNF4. (A) Left panel: SUMO conjugation assay with recombinant TOP1 incubated with SUMO E1, SUMO E2, SUMO-1, and increasing concentration of PIAS4. Reaction products were separated by SDS-PAGE and monitored by IB using anti-SUMO-1 antibody. Right panel: same as left panel except the IB was performed with anti-SUMO-2/3 antibody. (B)&(C) same as panel (A) except TOP2α and TOP2β were used instead of TOP1, respectively. (D) Ubiquitin conjugation assay with recombinant TOP1 and RNF4. Unmodified TOP1 (left) or *in vitro* SUMOylated TOP1 in the presence of SUMO E2 and PIAS4 was tested for ubiquitin conjugation in the presence of Ub E1, Ub E2 and the indicated concentrations of RNF4. Reaction products were separated by SDS-PAGE and monitored by IB using anti-Ub antibody. (E)&(F) same as panel (D) except TOP2α and TOP2β were used instead of TOP1, respectively.

**Figure S5.**
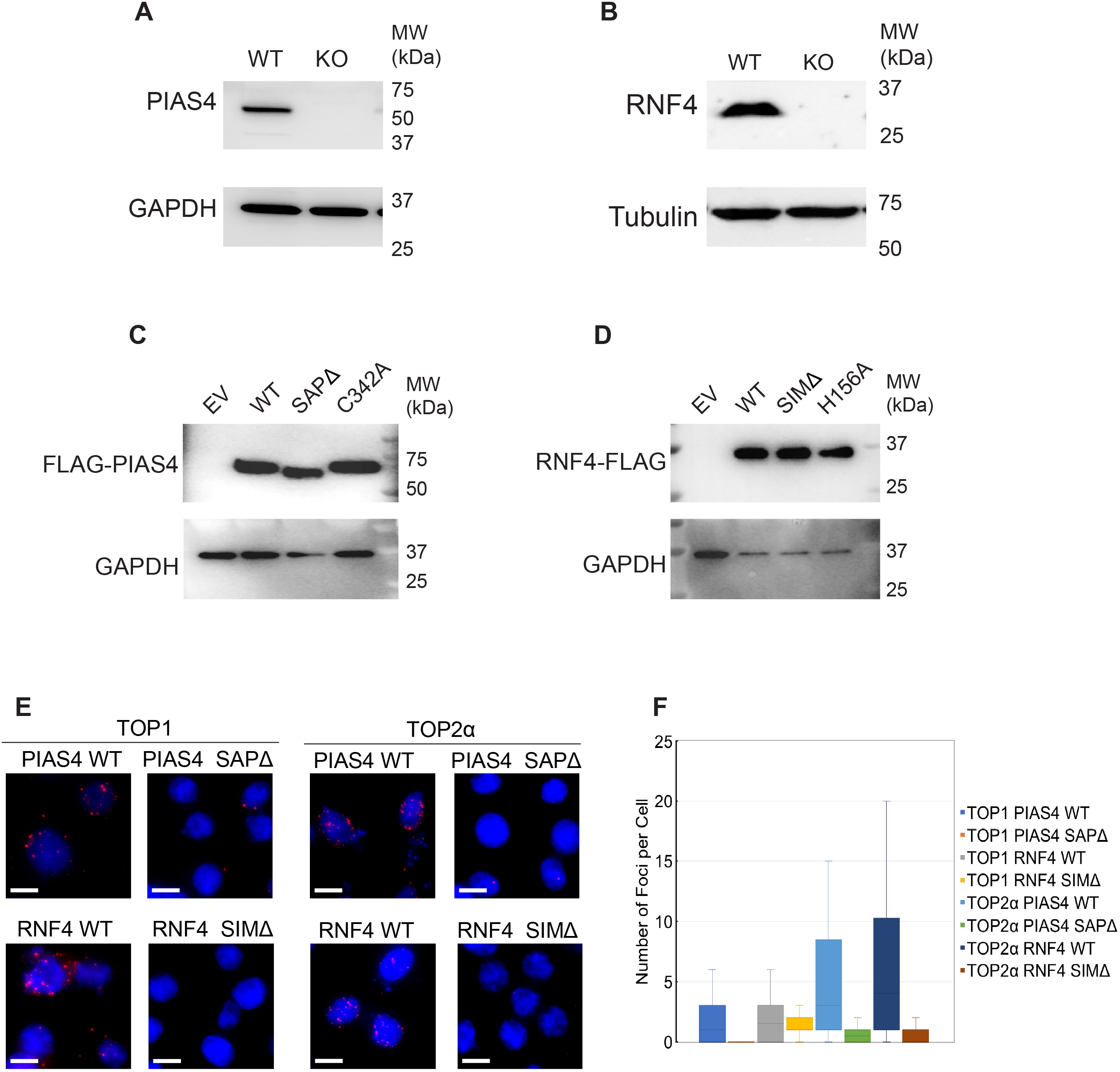
Coordination of PIAS4 and RNF4 for TOP-DPC repair. (A) WB of HCT116 WT and PIAS4 knockout (KO) cells using anti-PIAS4 antibody and anti-GAPDH antibody. (B) WB of MCF7 WT and RNF4 KO cells with or without RNF4-FLAG overexpression (OE) using anti-RNF4 antibody and anti-tubulin antibody. (C) HCT116 cells were transfected with empty vector (EV), FLAG-PIAS4 WT, FLAG-PIAS4, FLAG-PIAS4 SAPΔ or FLAG-PIAS4 C342A expression plasmid for 48 hours, WB using anti-FLAG antibody and anti-GAPDH antibody. (D) MCF7 cells were transfected with empty vector (EV) or RNF4-FLAG WT, RNF4-FLAG SIMΔ or RNF4-FLAG H156A expression plasmid for 48 hours, followed by WB using anti-FLAG antibody and anti-GAPDH antibody. (E) Left panel: U2OS cells were transfected with the indicated plasmids (FLAG-tagged) before CPT treatment (20 μM, 30 min), followed by PLA using anti-TOP1 and anti-FLAG antibodies. Right panel: U2OS cells were transfected with the indicated plasmids (FLAG-tagged) before ETP treatment (200 μM, 30 min), followed by PLA using anti-TOP2α and anti-FLAG antibodies. Scale bar represents 10 microns. (F) Quantitation of foci per cells of each treatment group as shown in panel (E).

**Figure S5.**
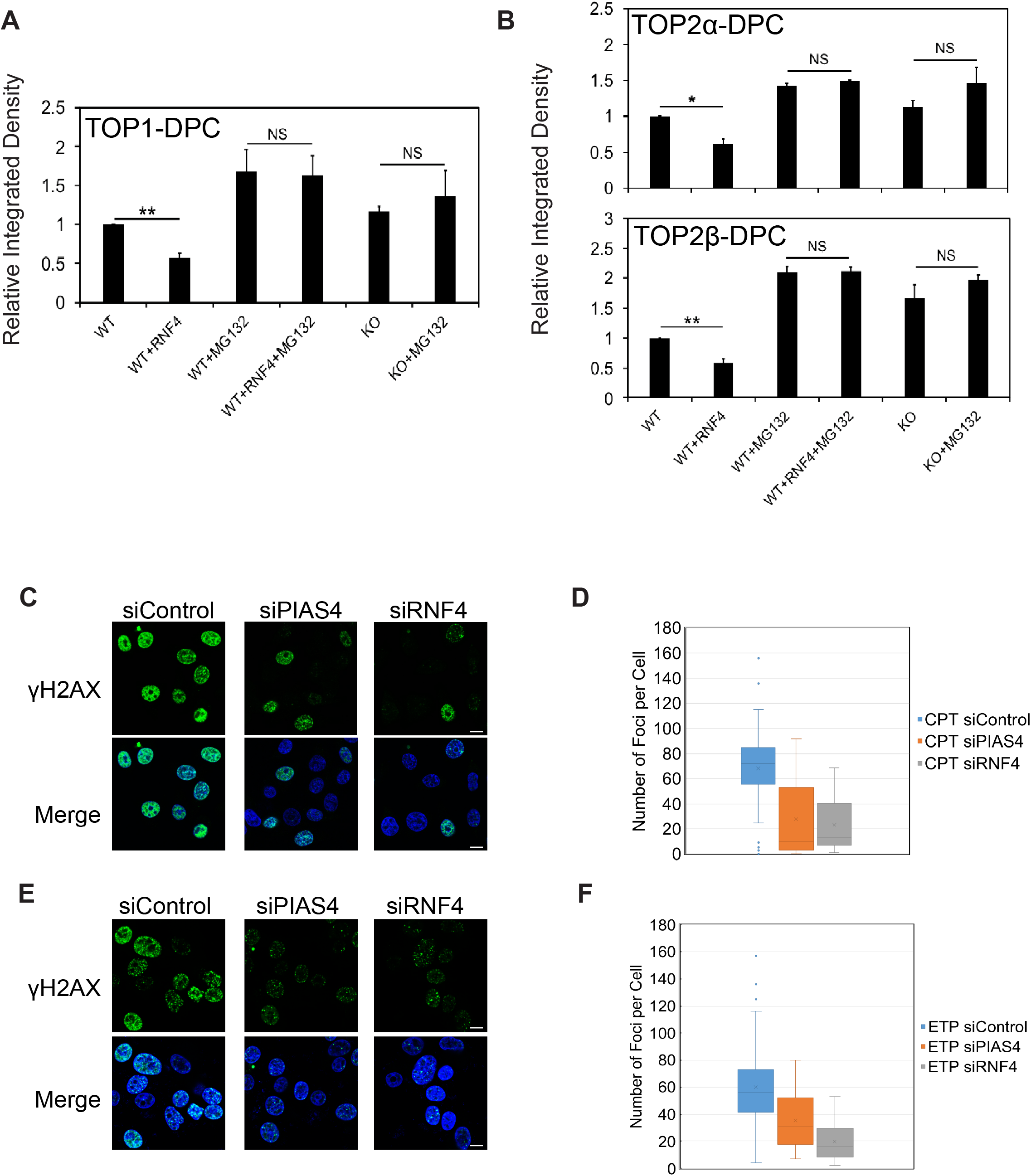
RNF4 Drives Proteasomal Degradation of TOP-DPCs. (A)&(B) Densitometric analyses comparing relative integrated densities of TOP1-, TOP2α- or TOP2β-DPC signals in independent experiments shown in Figure 7 panels (A)-(C). Integrated density of TOP-DPC signals of each group was normalized to MCF WT cells treated with CPT or ETP alone. * p < 0.05; ** p < 0.01. NS, not significant. (C) U2OS cells were transfected with the indicated siRNAs, followed by 1 h CPT (1 μM) treatment and then IF assay for detection of γH2AX. (D) Quantitation of foci per cells for each treatment group from experiments performed as shown in panel (C). Scale bar represents 20 microns. (E) U2OS cells were transfected with the indicated siRNAs, followed by 1h ETP (5 μM) treatment and IF assay for detection of γH2AX. (F) Quantitation of foci per cells of each treatment group from experiments performed as shown in panel (E). Scale bar represents 20 microns.

## MATERIALS AND METHODS

### Yeast Strains and Plasmids

BY4741 and YMM10 are two haploid strains of Saccharomyces cerevisiae, which were used as parental strains in this study. Individual BY4741 ORF KANMX4 deletion mutants (*slx5*Δ, *slx8*Δ, *siz1*Δ, *siz2*Δ, *mre11*Δ, *tdp1*Δ) were purchased from Open Biosystems or Dharmacon. To generate disruption of TDP1 in BY4741 *slx5Δ* and *siz1Δ* strains, plasmid containing TDP1 sequence was digested with restriction enzymes to create an internal deletion of the gene followed by replacement with a LEU2 cassette. Disrupted alleles were PCR amplified, and the PCR product was used in transformation of BY4741 *slx5Δ* and *siz1Δ* strains (Nitiss et al., 2006). To generate BY4741 *slx5Δsiz1Δ* double mutant, BY4742 *siz1Δ*::KANMX4:: and BY4741 *slx5Δ*::LEU2:: were mated, and diploid cells were sporulated and microdissected. Mating type was determined using H317 MATa and H318 MATα ura2 tester strains. In YMM10 strain, genes encoding nine different types of transmembrane drug-efflux pump proteins are deleted. A study has reported that YMM10 strain displays improved drug accumulation and enhanced sensitivity to ETP(Tombline et al., 2017). For immunodetection of yeast TOP1, N-terminally HA-tagged TOP1 cDNA was amplified by PCR using primers and inserted in NcoI/XhoI sites of pYX112 vector. Strains were transformed with pYX112 as an empty vector or pYX112 HA-TOP1 in which TOP1 expression is driven from the promoter TPI1. For immunodetection of yeast TOP2, strains were transformed with yCP50 as an empty vector or a yeast TOP2-3×HA overexpression plasmid pDED1 TOP2 in which TOP2 expression is driven from the DED1 promoter. To enhance accumulation of CPT and ETP in yeast cells, XhoI-excised construct of DNA binding domain (DBD) of PDR1 gene fused in frame to transcription repressor gene CYC8 from pBlueScript backbone was transformed to all BY4741 single mutant derivatives to repress Pdr1 regulated genes as described previously (Stepanov et al., 2008).

### Site-Directed Mutagenesis in Yeast Vectors

SUMOylation-deficient yeast TOP1 allele, *top1-K65, 91, 92R* (Chen et al., 2007) was constructed in pYX112 HA-TOP1 vector using oligonucleotides 5’-AGATTTCCAAGAAAAAGACTAAGAAAATAAGGACCGAACCAGTGCAA-3′(K65R), 5′-G GCGCGACATCAAAGCCTAAAAAAATCAGGAGAGAAGATGGTGATG-3′(K91,92R). SUMOylation-deficient yeast TOP2 allele, *top2-SNM* (Bachant et al., 2002) was constructed in pDED1 TOP2-3×HA vector using oligonucleotides 5’-GCTAGAAAGGGCAAAAAAATTAGGGTCGAGGATAAGAATTTTG-3′(K1220R), 5′-GCAAGGCGCCTACAAAGATTAGAAGAGAGAAAACGCCTTCTGTTTC-3′(K1246R, K1247R) and 5′-CTTCTTCTATTTTCGACATAAgGAgAGAAGATAAAGATGAGGGCG-3′(K1277R, K1278R).

### Western Blotting in Yeast

After inhibitor treatments, yeast cells were pelleted then washed with alkaline lysis buffer (200 mM NaOH, 2 mM EDTA) (Mao et al., 2000b). Cells were then resuspended in 700 μl alkaline lysis buffer and lysed by homogenization at 4 °C. After 4 cycles of homogenization (50 seconds each cycle, 5 min rest in between), lysates were centrifuged and 400 μl supernatants were retrieved, followed by neutralization by the addition of 80 μl of 1 M HCl, 600 mM Tris, pH 8.0. Samples were treated with 54 μl of 10× micrococcal nuclease buffer (50 mM CaCl2, 500 mM Tris pH=7.9) and 1 μl micrococcal nuclease (100 units/μl). The resulting mixtures were incubated on ice for 1h for releasing TOPs from DNA by digestion. Protein concentration were determined by using the Bradford assay (Bio-Rad). The lysates were mixed with 4× laemmli buffer and subjected to SDS-PAGE electrophoresis, followed by IB with indicated antibodies.

### Clonogenic Assay in Yeast

Drug sensitivity assay in yeast cells was carried out as described previously(Nitiss et al., 1992). Briefly, cells were grown to mid-exponential phase in synthetic complete media lacking uracil (SC-URA) then diluted to 2 × 10^6^ cells/ml. After addition of inhibitors, cells were incubated, diluted, and plated at various time points as indicated onto SC-URA plates. Plates were incubated at 30 °C, and the numbers of colonies were counted. Results were expressed relative to the number of viable colonies at the time of drug addition.

### HA-Immunoprecipitation in Yeast

Yeast lysates were prepared by the alkaline lysis procedure as described above. The lysates were incubated with anti-HA antibody in yeast IP lysis buffer containing 50 mM Tris-HCl, pH 7.5, 50 mM NaCl, 1% NP-40, 0.2% Trinton X-100, 5% glycerol, 1 mM DTT, 20 mM N-ethylmaleimide and yeast protease inhibitor cocktail (Sigma Aldrich, P8215) at 4 °C overnight. 50 μl Protein A/G PLUS-agarose slurry was added and incubated with the lysates for anther 4 hrs. After centrifugation, immunoprecipitates were washed with yeast IP lysis buffer 2 times then resuspended in 2 × laemmli buffer for SDS-PAGE and immunoblotting with indicated antibodies.

### ICE assay in Yeast

After inhibitor treatment, yeast cells were washed in lysis buffer containing 6 M guanidinium thiocyanate (GTC), 1% sarkosyl, 4% Triton X 100, 10 mM Tris–HCl (pH 7.0), 20 mM EDTA, yeast protease inhibitor cocktail, 20 mM N-ethylmaleimide and 1 mM DTT. Lysates were prepared in the lysis buffer by homogenization using a bead beater. Lysates were incubated at 65°C for 15 min, followed by centrifugation to remove cell debris. Supernatants were diluted in 1% sarkosyl in TE buffer and centrifuged again. Supernatants were then loaded on 150% (w/v) cesium chloride for ultracentrifugation for 18 hr at 42000 rpm in a NVT 65.2 rotor (Beckman coulter) at 25°C. Nucleic acids pellets containing TOP-DPCs were retrieved in ddH2O and digested with RNase A. Purified DNA samples were quantitated and 10 μg of each sample was digested with micrococcal nuclease and subjected to SDS-PAGE electrophoresis for immunodetection of total TOP-DPCs, ubiquitylated TOP-DPCs and SUMOylated TOP-DPCs. 2 μg of each sample was subjected to slot-blot for immunoblotting with anti-dsDNA antibody to confirm equal amounts of DNA load.

### Spot Test in Yeast

For yeast growth on media containing drug, 3 μl of a serial dilution of growing cells was applied to SC-URA plates containing the indicated concentrations of inhibitors. Plates were incubated at 30°C for 2 days and photographed.

### Human Cell Culture

HEK293 cells, HCT116 colon cancer cells, MCF7 breast cancer cells and HeLa cells were in cultured in DMEM medium (Life Technologies) supplemented with 10% (v/v) fetal bovine serum, 100 units/ml penicillin, 100 μg streptomycin /ml streptomycin and 1x GlutaMax in tissue culture dishes at 37 °C in a humidified CO2 – regulated (5%) incubator.

### Generation of Gene Knock-out Cells Using CRISPR-Cas9

To stably knockout the genes encoding TOP2B, PIAS4 or RNF4, the CRISPR–Cas9 genome-editing method was used(Ran et al., 2013). To delete TOP2B in HeLa cells, two 25-bp (minus the PAMs) guide RNA sequences 5’-CACCGGATTTGGCTGGTTCGTGTAG-3’ and 5’-ACCGTAAATTTGGACAGATCTGGT-3’ targeting TOP2B exon 7 were designed using the CHOP CHOP tool and cloned into the Cas9 expressing guide RNA vectors pX458 and pX459, respectively. In brief, the Bbs1 cutting site containing guide RNA sequences were annealed and then cloned into the guide RNA vector using T4 ligase (New England Biolabs). The plasmids were transfected with Lipofectamine 3000 in HeLa cells. Transfected cells were enriched by selection in 0.5 mg/ml puromycin containing media for 3 days prior to isolation of single clones and screening for loss of TOP2β by Western blotting. To delete PIAS4 in HCT116 cells, two guide RNA sequences 5’-CACCGCATGCACTCCACCTACGACC-3’ and 5’-CACCGAGAACGGCAGCTTCACCAGG-3’ targeting PIAS4 exon 2 were designed and cloned into pX458 and pX459, respectively. The plasmids were transfected in HCT116 cells, followed by selection in media containing 1 mg/ml puromycin for 3 days prior to isolation of single clones and screening for loss of PIAS4 by Western blotting. To delete RNF4 gene in MCF7 cells, two guide RNA sequences 5’-CACCGAGGCAAAGAAAATCGAGACC-3’ and 5’-CACCGCATACACTCTCGTCCACGGC-3’ targeting RNF4 exon 3 were designed and cloned into pX458 and pX459, respectively. The RNF4 guide constructs were transfected into MCF7 cells, followed by puromycin selection and single clone isolation. Deletion of RNF4 gene was confirmed by sanger sequencing and loss of RNF4 expression by Western blotting.

### Expression Plasmids and siRNAs in Human Cells

For human TOP1 expression in HEK293 cells, N-terminally 6×His tagged TOP1 cDNA was amplified by PCR using primers 5’-GCATATCCTCGAGCGCCACCATGGCCCATCATCACCATCACCACAGTGGGGACCACCTCC ACAAC −3’ and 5’-GCTAGAGCGGCCGCCTAAAACTCATAGTCTTCATCAGC −3’ and using pGALhTOP1 yeast plasmid as template. The PCR product was inserted into PspXI/NotI sites of a pT-REx-DEST Gateway vector (Invitrogen). For human TOP2α expression in HEK293 cells, N-terminally FLAG-tagged TOP2α cDNA was amplified by PCR using primers 5’-GCATATCCTCGAGCGCCACCATGGACTACAAGGACGACGATGACAAGGAAGTGTCACCATTGCAGCCTGTAAAT-3’ and 5’-GCTAGAGCGGCCGCTTAAAACAGATCATCTTCATCTGACTC-3’ and using pMJ1hTOP2α yeast plasmid as template then inserted into PspXI/NotI sites of the pT-REx-DEST Gateway vector. N-terminally FLAG-tagged TOP2β cDNA (derived from pHT500hTOP2β yeast plasmid) was PCR-amplified using primers 5’-GCATATCCTCGAGCGCCACCATGGACTACAAGGACGACGATGACAAGGCCAAGTCGGGTGGCTGCGGCGCGGGA-3’ and 5’-GCATATCCTCGAGCGCCACCATGGCCCATCATCACCATCACCACGCCAAGTCGGGTGGCTGCGGCGCGGGA-3’ and inserted into PspXI/NotI sites of the pT-REx-DEST Gateway vector. HA-Ubiquitin WT plasmid was a gift from Ted Dawson (Addgene plasmid # 17608). HA-SUMO-1 WT plasmid was a gift from Guy Salvesen (Addgene plasmid # 48966). HA-SUMO-2 WT plasmid was a gift from Guy Salvesen (Addgene plasmid # 48967). Flag-PIAS4 WT plasmid was a gift from Ke Shuai (Addgene plasmid # 15208). RNF4-Myc-FLAG WT plasmid was purchased from OriGene (CAT#: RC207273). PSMB5-Myc-FLAG WT plasmid was purchased from OriGene (CAT#: RC209326L3). siRNA transfections were performed using Lipofectamine RNAiMAX (Invitrogen) according to the manufacturer’s instructions. All siRNAs were used at a final concentration of 50 nM unless otherwise indicated. The following siRNAs were used: control siRNA (Dharmacon, CAT# D-001206-13-05); TOP2A siRNA (Dharmacon, CAT# M-004239-00-0005); SUMO-1 siRNA (Dharmacon, CAT# M-016005-00-0005); SUMO-2 siRNA (Dharmacon, CAT# L-016450-00); SUMO-3 siRNA (Dharmacon, CAT# L-019730-00); UBC9 siRNA (Dharmacon, CAT# M-004910-00-0005); TOPORS siRNA (Dharmacon, CAT# M-020048-00-0005); PIAS1 siRNA (Dharmacon, CAT# M-008167-00-0005); PIAS4 siRNF1 (Dharmacon, CAT# M-006445-00-0005 RNF4 siRNA (Dharmacon, CAT# M-006557-00-0005), RNF111 siRNA (Dharmacon, CAT# M-007002-00-0005).

### Site-Directed Mutagenesis (SDM) in Mammalian Expression Vectors

pTrex-6×His-TOP1Y723F was generated by QuikChange II XL SDM kit (Agilent) using oligonucleotides 5’-CTAGGGTCCAGAAAATTGAGTTTGGAGGTTCCCAGG-3’ pTrex-6×His-TOP1 K117, K153, K103R (CPT3KR) was generated using oligonucleotides 5’-TCTTCAGGTTCATCTCTAATTTGTGGTGGACTAGAGAAGC-3’ (K117R), 5’-TCTTGGTATCTTCTGTTCTAATTTTCTTAGGTTTATAATCAGCATCATCC-3’ (K153R), 5’-TTCCTTCTCCTTCCTTATTTTTGCATCCCCAGAGGCT-3’ (K103R). Ubiquitin K6R was generated by Q5 SDM Kit using oligonucleotide 5’-ATCTTCGTGAGGACCCTGACTGG-3’. Ubiquitin K11R was generated using oligonucleotide 5’-CTGACTGGTAGGACCATCACTC-3’. Ubiquitin K27R was generated using oligonucleotide 5’-GAGAATGTCAGGGCAAAGATCC-3’. Ubiquitin K29R was generated using oligonucleotide 5’ GTCAAGGCAAGGATCCAAGAC-3’. Ubiquitin K33R was generated using oligonucleotide 5’-ATCCAAGACAGGGAAGGCATC-3’. Ubiquitin K48R was generated using oligonucleotide 5’-TTTGCTGGGAGACAGCTGGAA-3’. Ubiquitin K63R was generated using oligonucleotide 5’-AACATCCAGAGAGAGTCCACCC-3’. HA-SUMO-1 K7R was generated using oligonucleotide 5’-CAGGAGGCAAGACCTTCAACTG-3’. HA-SUMO-2 K11R was generated using oligonucleotide 5’-GAAGGAGTCAGGACTGAGAACAAC-3’. FLAG-PIAS4 C342A was generated using oligonucleotide 5’-GGCAGAGACCGCCGCCCACCTGCAG-3’. FLAG-PIAS4 SAPΔ was generated using oligonucleotide 5’-CAGTTTGACTGTAGCCCTG-3’. RNF4 SIM2, 3, 4 Δ-FLAG was generated using oligonucleotides 5’-GACGCCACTTGTGAATCTTTAGAGC-3’ (SIM2Δ), 5’-GATGCGACTCACAATGACTCTG-3’ (SIM3Δ), 5’-GCTGCAGACGAAAGAAGAAGACCAAG-3’ (SIM4Δ). RNF4 H156A-FLAG was generated using oligonucleotide 5’-AGAATGCGGCGCTGTCTTCTGTAGC-3’.

### Western Blotting in Human Cells

CPT-induced TOP1 degradation and ETP-induced TOP2 degradation were monitored by Western blotting of the alkaline lysates prepared from drug-treated HEK293 cells with slight modifications(Ban et al., 2013). Following treatment, cells were washed with DMEM and incubated at 37°C in a CO_2_ incubator for 30 min then lysed with 100 μl of an alkaline lysis buffer (200 mM NaOH, 2 mM EDTA). Alkaline lysates were neutralized by the addition of 100 μl of 1 M HEPEs buffer, pH 7.3, followed by mixing with 10 μl 100mM CaCl_2_, 1 μl 2M DTT and 2 μl 100 × protease inhibitor cocktail and 200 units of micrococcal nuclease. The resulting mixtures were incubated on ice for 1 h. 70 μl of 4 × laemmli buffer was added to each sample. The lysates were boiled for 10 min, analyzed by SDS-PAGE, and immunoblotted with various antibodies as indicated. Other proteins were detected by lysing cells with RIPA buffer (150mM NaCl, 1% NP-40, 0.5% Sodium deoxycholate, 0.1% SDS, 50 mM Tris pH 7.5, 1 mM DTT and protease inhibitor cocktail).

### In vivo of complex (ICE) assay in Human Cells

TOP-DPCs were isolated and detected using in vivo complex of enzyme (ICE) assay as previously described(Anand et al., 2018). Briefly, cells were lysed in sarkosyl solution (1% w/v) after treatment. Cell lysates were sheared through a 25g 5/8 needle (10 strokes) to reduce the viscosity of DNA and layered onto CsCl solution (150% w/v), followed by centrifugation in NVT 65.2 rotor (Beckman coulter) at 42,000 RPM for 20 hours at 25 °C. The resulting pellet containing nucleic acids and TOP-DPCs was obtained and dissolved in TE buffer. The samples were quantitated and subjected to slot-blot for immunoblotting with various antibodies as indicated. 2 μg of DNA is applied per sample. For mass spectrometric analysis, ICE samples were treated with RNase A to eliminate RNA contamination. TOP-DPCs were quantified by densitometric analysis using ImageJ.

### Detection of Ubiquitylated and SUMOylated Topoisomerase DNA-Protein Crosslinks (DUST) Assay in Human Cells

After topoisomerase inhibitor treatments, 1 × 10^6^ human cells in 35 mm dish per sample were washed with 1 × PBS and lysed with 600 μl DNAzol (Invitrogen), followed by precipitation with 300 μl 200 proof ethanol. The nucleic acids were collected, washed with 75% ethanol, resuspended in 200 μl TE buffer then heated at 65°C for 15 minutes, followed by shearing with sonication (40% output for 10 sec pulse and 10 sec rest for 4 times). The samples are centrifuged at 15,000 rpm for 5 min and the supernatant are collected and treated RNase A (100 μg/ml) for 1 h, followed by addition of 1/10 volume of 3 M sodium acetate sodium acetate and 2.5 volume of 200 proof ethanol. After 20 min full speed centrifugation, DNA pellet was retrieved and resuspended in 100 μl TE buffer for spectrophotometric measurement to quantitate DNA content. 10 μg of each sample was digested with 50 units micrococcal nuclease (Thermo Fisher Scientific, 100 units/μl) in presence of 5 mM CaCl_2_, followed by SDS-PAGE electrophoresis for immunodetection of total TOP-DPCs, SUMOylated TOP-DPCs as well as ubiquitylated TOP-DPCs using specific antibodies. In addition, 2 μg of each sample was subjected to slot-blot for immunoblotting with anti-dsDNA antibody (Abcam, ab27156) to confirm equal DNA loading.

### His-Pull Down Assay in Denaturing and Native conditions

Human cells are washed with 1 × PBS and incubated with 220 μl IP lysis buffer (5 mM Tris-HCl pH 7.4, 150 mM NaCl, 1 mM EDTA, 1% NP-40, 0.2% Triton X-100, 5% glycerol, 1 mM DTT, 20 mM N-ethylmaleimide and protease inhibitor cocktail) on a shaker for 15 min at 4 °C, followed by sonication and centrifugation. Supernatant was collected and treated with 1 μl benzonase (250 units/μl) for 1h. An aliquot (20 μl) of the lysate of each treatment group was saved as input. The rest of the lysates was divided in two groups: native pull-down and denaturing pull-down. For native pull-down, lysates were resuspended in 900 μl IP lysis buffer containing 10 mM imidazole and 100 μl equilibrated Ni-NTA-agarose and rotated overnight at 4°C. Ni-NTA resin was spun down and washed with TI buffer two times (25 mM Tris HCL, 20 mM imidazole, pH 6.8), followed by resuspension in 2 × laemmli buffer for SDS-PAGE and immunoblotting with various antibodies as indicated. For denaturing pull-down, lysates were resuspended in 900 μl Buffer A (6M guanidine-HCL, 0.1 M Na_2_HPO4/NaH_2_PO4, 10 mM imidazole pH8.0) containing 100 μl equilibrated Ni-NTA-agarose and rotated overnight at 4°C. Ni-NTA resin was spun down and washed with TI buffer two times (25 mM Tris HCL, 20 mM imidazole, pH 6.8), followed by resuspension in 2 × laemmli buffer for SDS-PAGE and immunoblotting with various antibodies as indicated.

### FLAG Immunoprecipitation (IP) in Denaturing and Native Conditions

Human cells are washed with 1 × PBS and incubated with 220 μl IP lysis buffer (5 mM Tris-HCl pH 7.4, 150 mM NaCl, 1 mM EDTA, 1% NP-40, 0.2% Triton X-100, 5% glycerol, 1 mM DTT, 20 mM N-ethylmaleimide and protease inhibitor cocktail) on a shaker for 15 min at 4 °C, followed by sonication and centrifugation. Supernatant was collected and treated with 1 μl benzonase (250 units/μl) for 1h. An aliquot (20 μl) of the lysate of each treatment group was saved as input. The rest of the lysates was divided in two groups: native IP and denaturing IP. For native IP, lysates were resuspended in 900 μl IP lysis buffer containing 2.5 μl anti-FLAG M2 antibody and rotated overnight at 4°C. 50 μl Protein A/G PLUS-agarose slurry was added and incubated with the lysates for anther 4 hrs. After centrifugation, immunoprecipitates were washed with RIPA buffer 2 times then resuspended in 2 × laemmli buffer for SDS-PAGE and immunoblotting with various antibodies as indicated. For denaturing pull-down, lysates were resuspended in 900 μl RIPA buffer containing 2.5 μl anti-FLAG M2 antibody and rotated overnight at 4°C. 50 μl Protein A/G PLUS-agarose slurry was added and incubated with the lysates for anther 4 hrs. After centrifugation, immunoprecipitates were washed with RIPA buffer 2 times then resuspended in 2 × laemmli buffer for SDS-PAGE and immunoblotting with various antibodies as indicated.

### Mass Spectrometry

Samples were either separated by SDS-PAGE for in-gel trypsin digestion as described(Shevchenko et al., 2006) or in-solution digested with trypsin following the filter-aided sample preparation (FASP) protocol as previously described(Wisniewski et al., 2009). Dried peptides were solubilized in 2% acetonitrile, 0.5% acetic acid, 97.5% water for mass spectrometry analysis. They were trapped on a trapping column and separated on a 75 μm x 15 cm, 2 μm Acclaim PepMap reverse phase column (Thermo Scientific) using an UltiMate 3000 RSLCnano HPLC (Thermo Scientific). Peptides were separated at a flow rate of 300 nL/min followed by online analysis by tandem mass spectrometry using a Thermo Orbitrap Fusion mass spectrometer. Peptides were eluted into the mass spectrometer using a linear gradient from 96% mobile phase A (0.1% formic acid in water) to 55% mobile phase B (0.1% formic acid in acetonitrile). Parent full-scan mass spectra were collected in the Orbitrap mass analyzer set to acquire data at 120,000 FWHM resolution; ions were then isolated in the quadrupole mass filter, fragmented within the HCD cell (HCD normalized energy 32%, stepped ± 3%), and the product ions analyzed in the ion trap. Proteome Discoverer 2.2 (Thermo) was used to search the data against human proteins from the UniProt database using SequestHT. The search was limited to tryptic peptides, with maximally two missed cleavages allowed. Cysteine carbamidomethylation was set as a fixed modification, and methionine oxidation set as a variable modification. Diglycine modification to lysine was set as a variable modification for experiments to identify sites of enzymatic PTMs. The precursor mass tolerance was 10 ppm, and the fragment mass tolerance was 0.6 Da. The Percolator node was used to score and rank peptide matches using a 1% false discovery rate.

### Recombinant Proteins

Human TOP1 was purified from baculovirus as previously described(Laco and Pommier, 2008). Human TOP2α and β were purified from yeast strains JEL1 *top1Δ* transformed with 12-URA-B 6×His-hTOP2α and JEL1 *top1Δ* transformed with 12-URA-C 6×His-hTOP2β. Induction of TOP2 by galactose as described previously(Dong et al., 2000). Yeast cells were lysed in equilibration buffer (300 mM KCl, 10 mM imidazole, 20mM Tris HCl pH 7.7, 10% glycerol and protease inhibitors) by glass bead homogenization. Lysates were incubated with Ni-NTA resin and washed using wash buffer #1 (300 mM KCL, 30 mM imidazole, 20M Tris HCl pH 7.7, 10% glycerol and protease inhibitors) then wash buffer #2 (150 mM KCl, 30 mM imidazole, 20 mM Tris HCl pH 7.7, 10% glycerol and protease inhibitors). hTOP2α and β were eluted on a Poly-Prep chromatography columns (Bio-Rad) with elution buffer (150 mM KCl, 300 mM imidazole, 20 mM Tris HCl pH 7.7, 10% glycerol and protease inhibitors). The peak protein fractions were dialyzed in dialysis buffer (750mM KCl, 50 mM Tris HCl pH 7.7, 20% glycerol, 0.1 mM EDTA and 0.5mM DTT) and His tag was removed using TEV protease. Recombinant human PIAS4 was purchased from OriGene (Cat. # TP306748). Recombinant human SUMO-1 and SUMO-2 were purchased from R&D systems (Cat. # K-700). Recombinant human SUMO activating enzyme E1 (SAE1/UBA2) was purchased from R&D systems (Cat. # E-315) Recombinant Human Ubc9 was purchased from (Cat. # E2-645). Recombinant human RNF4 was purchased from R&D Systems (Cat. # E3-210-050). Recombinant human ubiquitin was purchased from R&D Systems (Cat. # U-100H). Recombinant human ubiquitin-activating enzyme E1 (UBE1) was purchased from R&D systems (Cat. # E-305). Recombinant human UbcH5a was purchased from R&D systems (Cat. # E2-616).

### In vitro SUMO Conjugation Assay

10 μl in vitro SUMOylation assay reaction in 1 x SUMO conjugation reaction buffer (R&D systems Cat. # B-60) contains 10 mM Mg^2+^-ATP solution pH 7.0 (R&D systems Cat. # B-20), protease inhibitor cocktail, 100 nM TOP1, TOP2α or TOP2β, 10 μM SUMO-1 or SUMO-2, 50 nM SUMO E1, 0.1 μM Ubc9 and PIAS4 of indicated concentrations. Reactions were incubated at 37 °C for 30 minutes, followed by SDS-PAGE and immunoblotting with anti-SUMO-1 (abcam ab32058) or anit-SUMO-2/3 antibodies.

### In vitro Ubiquitin Conjugation Assay

10 μl in vitro ubiquitylation assay reactions in 1 x ubiquitin conjugation reaction buffer (R&D systems Cat. # B-70) contains 10 mM Mg^2+^-ATP solution pH 7.0 (R&D systems Cat. # B-20), protease inhibitor cocktail, 100 nM TOP1, TOP2α or TOP2β, 10 μM ubiquitin, 50 nM ubiquitin E1, 0.1 μM UbcH5a and RNF4 of indicated concentrations. Reactions were incubated at 37 °C for 30 minutes, followed by SDS-PAGE and immunoblotting with anti-ubiquitin antibody (Santa Cruz sc-8017).

### In vitro SUMOylation-Ubiquitylation Coupled Assay

In vitro SUMOylation assay reaction was conducted in 1 x SUMO conjugation reaction buffer containing 350 nM TOP1, TOP2α or TOP2β, 10 μM SUMO-1 or SUMO-2, 50 nM SUMO E1, in presence or absence of 0.1 μM Ubc9 and 0.5 μM PIAS4 at 37 °C for 30 min. The reaction was then aliquoted and diluted in 1 x ubiquitin conjugation reaction buffer containing 10 mM Mg^2+^-ATP solution pH 7.0, protease inhibitor cocktail, 10 μM ubiquitin, 50 nM ubiquitin E1, 0.1 μM UbcH5a and RNF4 of indicated concentrations. Reactions were incubated at 37 °C for 30 minutes, followed by SDS-PAGE and immunoblotting with anti-ubiquitin antibody.

### TOP1-DPC Immunofluorescence

TOP1-DPC immunofluorescence was performed using previously described protocol(Patel et al., 2016) with slight modification. U2OS cells grown on chamber slides were treated with CPT in absence or presence of indicated inhibitors. After inhibitor treatments, cells were washed with PBS and fixed for 15 min at 4°C in 4% paraformaldehyde in PBS and permeabilized with 0.25% Triton X-100 in PBS for 15 min at 4°C. The samples were incubated in 2% SDS at room temperature for 10 min, washed and blocked with PBS containing 0.01% Triton X-100, 0.05% Tween 20 and 1% bovine serum albumin (PBSTT-1%BSA). After reaction overnight with TOP1-DPC antibody (Millipore Sigma) in PBSTT-BSA at 4°C, cells were rinsed with PBSTT and incubated with Alexa Fluor 568-conjugated secondary antibody (Invitrogen) at 1:1000 in PBSTT-BSA for 1 h in subdued light; washed and mounted using mounting medium with DAPI (Vectashiled). Images were captured on an instant structured illumination microscope, processed using ImageJ and analyzed using Imaris.

### γH2AX Immunofluorescence

U2OS cells were seeded on chamber slides. After inhibitor treatment, cells were washed with PBS and fixed for 15 min at 4°C in 4% paraformaldehyde in PBS and permeabilized with 0.25% Triton X-100 in PBS for 15 min at 4°C. The samples were blocked with PBSTT-1% BSA, followed by overnight incubation with γH2AX antibody (Milipore Sigma) in PBSTT-BSA at 4°C, cells were rinsed with PBSTT and incubated with Alexa Fluor 4888-conjugated secondary antibody (Invitrogen) at 1:1000 in PBSTT-BSA for 1 h in subdued light; washed and mounted using mounting medium with DAPI (Vectashiled). Images were captured on Zeiss LSM 880/Airyscan confocal microscope and processed using ImageJ.

### Proximity Ligation Assay (PLA)

Duolink PLA fluorescence assay (Sigma Aldrich) was performed following manufacturer’s instruction. In brief, U2OS cells were seeded on coverslips and treated with CPT or ETP. After inhibitor treatment, cells were washed with PBS and fixed for 15 min at 4°C in 4% paraformaldehyde in PBS and permeabilized with 0.25% Triton X-100 in PBS for 15 min at 4°C. The coverslips were blocked with Duolink blocking solution and incubated with indicated antibodies in the Duolink antibody diluent overnight, followed by incubation with PLUS and MINUS PLA probes, ligation and amplification. Coverslips were then washed and mounted with using mounting medium with DAPI. Images were captured on wide field microscope, processed using ImageJ and analyzed using Imaris.

### Statistical Analyses

Error bars on bar graphs represent standard deviation (SD) and p-value was calculated using paired student’s t-test for independent samples.

